# Joint cmICA: auto-linking of structural and functional connectivity

**DOI:** 10.1101/2022.09.12.507415

**Authors:** Lei Wu, Vince Calhoun

## Abstract

The study of human brain connectivity, including structural connectivity (SC) and functional connectivity (FC), provides insights into the neurophysiological mechanism of brain function and its relationship to human behavior and cognition. Both types of connectivity measurements provide crucial yet complementary information. However, integrating these two modalities into a single framework remains a challenge, because of the differences in their quantitative interdependencies as well as their anatomical representations due to distinctive imaging mechanisms. In this study, we introduced a new method, joint cmICA (connectivity matrix ICA), which provides a data-driven parcellation and automated-linking of SC and FC information simultaneously using a joint analysis of functional MRI and diffusion-weighted MRI data. We showed that these two connectivity modalities produce common cortical segregation, though with various degrees of (dis)similarity. Moreover, we show conjoint functional connectivity networks and structural white matter tracts that directly link these cortical parcellations/sources, within one analysis. Overall, data driven joint cmICA provides a new approach for integrating or fusing structural connectivity and functional connectivity systematically and conveniently, and provides an effective tool for connectivity-based multimodal data fusion in brain.

## Introduction

Understanding neural network organizations in the human brain is one of the primary goals in neuroscience. The complexity of brain computation mechanism implies that it is a product of combination of local synchronizations as well as longer ranged connections between neurons (Jbabdi & Behrens, 2013). The term ‘brain connectivity’ in neuroscience typically refers to a systematical pattern of structural/anatomical linkages and/or functional interactions between different neuronal nodes within one brain nervous system (E. Allen, Damaraju, Eichele, Wu, & Calhoun, 2018; Elena A. Allen et al., 2012; V. D. Calhoun, Miller, Pearlson, & Adalı, 2014; Karl J Friston, 2011; Wu, Calhoun, Jung, & Caprihan, 2015; Wu, Caprihan, Bustillo, Mayer, & Calhoun, 2018). The nodes here can be designated as segregated brain parcels, neuronal populations, or even single neurons, depending on the data. And a more comprehensive investigation of these linkages and interactions can benefit from data driven approaches which can identify complex spatial and temporal relationships such as interacting brain networks.

Recent focus on multimodal neuroimaging combining functional MRI (fMRI) and diffusion MRI (dMRI) has led to a better understanding of brain communication and its regulation of functional integration and segregation (Chu, Parhi, & Lenglet, 2018; Deco, Tononi, Boly, & Kringelbach, 2015; Hagmann et al., 2008). White matter tracts are the neuroanatomical pathways that directly connect functional network regions and facilitate the travel of interregional information (Honey et al., 2009). Multiple brain disorder studies, e.g., Alzheimer’s disease and dementia, also suggest that a subtle change in white matter connectivity may contribute greatly to the impairment of functional networks that are associated with brain cognitive decline (Taylor et al., 2017). Therefore, methods that provide improved mapping of structural connectivity architecture and functionally linked brain networks using whole-brain modeling (Deco et al., 2015), may play a vital role in discovering underlying brain mechanisms.

Numerous methods have been developed to incorporate dMRI and fMRI data together, which utilizes the complementary information provided from both modalities. However, most approaches process the dMRI information separately and sometimes use it to constrain the fMRI information (Crimi, Dodero, Sambataro, Murino, & Sona, 2021; Deco, Kringelbach, Jirsa, & Ritter, 2017; Sihag et al., 2022). Other fusion approaches use seeds or regions of interests (ROIs) from one or both modalities to associate SC and FC (Bonnelle et al., 2012; Tarun, Behjat, Bolton, Abramian, & Van De Ville, 2020). Some data-driven fusion approaches such as joint independent component analysis (jICA) (V. D. Calhoun & Sui, 2016; Franco et al., 2008; Teipel et al., 2010), multiple-set canonical correlation Analysis (MCCA) (Fu et al., 2020; Sui et al., 2015) and their extensions have been applied to derive spatial decompositions from two modalities based on inter-subject covariation, but they usually require summarizing modalities into features first. Thus, there is still a need for data fusion approaches which can 1) provide data driven parcellations, 2) automatically capture spatial correspondence among the structural and functional parcellations to fully leverage the information within each image type, and 3) address limitations of current approaches with a goal of providing more flexibility in estimating sources and connectivity maps. The proposed joint cmICA provides a natural extension of existing work in this regard.

The connectivity matrix independent component analysis (cmICA) method (Wu et al., 2015) was developed to extract maximally independent spatial sources and their corresponding connectivity maps to the whole brain via the estimation of a connectivity matrix at the group level. cmICA was initially developed to decompose the tractographic connectivity matrix from dMRI data (Wu et al., 2015; Wu, Caprihan, & Calhoun, 2017b), and found to be particularly sensitive to subtle connectivity changes across subjects (Elena A Allen et al., 2011; Koch et al., 2010; Wu et al., 2015), longer-range inter-regional connectivity and spatial localization compared to conventional ICA methods (Wu et al., 2015). Following the initial approach, we have since extended and optimized cmICA from structural to functional connectivity and dynamic functional connectivity using BOLD data (Wu et al., 2018; Wu, Caprihan, & Calhoun, 2017a; Wu, Caprihan, & Calhoun, 2021). In this study, we further extend cmICA (Wu et al., 2018) to jointly extract cortical source segregations from both structural and functional connectivity modalities, and to link the corresponding functional networks and structural networks (i.e., fiber tracts) if existing and well formed.

## Material and Methods

### 1. Subjects

Subjects were recruited via a Center Of Biomedical Research Excellence (COBRE, http://cobre.mrn.org) program at the Mind Research Network (MRN). In this paper, we used Fmri and diffusion MRI (dMRI) data from the control subjects only (n = 60, age = 36.8 ± 12.1 years) which we previously studied to estimate structural and functional connectivity separately in unimodal cmICA analyses (Wu et al., 2015; Wu et al., 2018). In this case we exclude the patients to focus on general connectivity pattern extraction as well as 1 HC subject that did not have both imaging modalities collected. All the participants were enrolled into the COINS platform (http://coins.trendscenter.org) (Scott et al., 2011; Wood et al., 2015). Prior to inclusion in the study, subjects were screened to ensure they were free from neurological or psychiatric disorders (DSM-IV Axis I) as well as active substance use disorders (except for nicotine).

### 2. Data acquisition and preprocessing

Data were collected and preprocessed as described in (Wu et al., 2015; Wu et al., 2018), with additional graphics processing unit (GPU) computational upgrades. All participants were scanned on a 3Tesla Siemens TIM Trio equipped with a 12-channel radio frequency coil. Subjects were instructed to fixate on a central crosshair presented with eyes open at rest.

Diffusion data were acquired via a single-shot spin-echo echo planar imaging (EPI) with a twice-refocused balanced echo sequence to reduce eddy current distortions. The dMRI sequence included 30 directions, b=800 s/mm^2^ and 5 measurements of b=0, for 6 minutes of acquisition time. The field of view (FOV) was 256 × 256 mm with a 2 mm slice thickness, 72 slices, 128 × 128 matrix size, voxel size = 2 mm x 2 mm x 2 mm, TE=84ms, TR=9000ms, number of excitations (NEX)=1, partial Fourier encoding of 3/4, and with a GRAPPA acceleration factor of 2. The analysis was performed using the FSL software package (http://fsl.fmrib.ox.ac.uk). Motion and eddy current correction were applied; gradient directions with signal dropout caused by subject motion were not included in further analysis; bedpostx GPU and probtrackx GPU were applied for ODF diffusion parameter estimation and probabilistic tractography. We used individual voxels over the entire brain as the seed regions as well as target regions, without pre-defining any ROI masks to provides data driven blind source separation. A 5mm spatial resolution and 2000 streamlines from each seed voxel were used to balance the computational speed and performance. Down-sampling at this stage also reduced the number of voxels and the connectivity matrix to a large, yet manageable size. After probabilistic tractography, whole brain voxel-paired connectivity profiles were converted to a two-dimensional connectivity matrix, by estimating the fiber counts between each voxel pair within their corresponding matric elements.

Resting state functional scans were collected by means of T2*-weighted gradient echo planar imaging sequences at 5 minutes long 150 volumes with the following parameters: repeat time (TR) = 2s, echo time (TE) = 29ms, field of view = 240mm, acquisition matrix = 64×64, flip angle = 75°, voxel size = 3.75×3.75×4.55mm^3^, gap = 1.05mm, 33 slices, ascending acquisition. A high-resolution structural anatomy was also acquired via a 3D MPRAGE T1 sequence (sagittal; matrix, 256×256; FOV, 256 mm; slice thickness, 1 mm; no gap; in-plane voxel size, 1 mm × 1 mm; flip angle, 7°; TR, 2.53 s; TE, 1.64 ms) to provide the anatomical reference for the functional scan. FMRI data were preprocessed using the software package SPM. The first five volumes were discarded to allow for T1 equilibration. Images were then realigned using the INRIalign motion correction algorithm (Freire & Mangin, 2001). Slice timing correction was applied using the middle slice as the reference frame. Next, data were spatially normalized into standard Montreal Neurological Institute space (K. J. Friston et al., 1995), spatially smoothed with a Gaussian kernel of FWHM 5mm and sub-sampled to 5×5×5 mm^3^, to match the spatial resolution and geometry of the dMRI data and reduce computational complexity. In addition, the data were detrended, despiked and temporal band-pass filtered from 0.001 to 0.15 Hz, to remove noise sources e.g., scanner drift, motion spikes and other non-specific high frequency artifacts (Elena A Allen et al., 2011; Cordes et al., 2000; Wu et al., 2018). In this study, a white matter mask and a gray matter mask (FSL) were used for more accurate cortical parcellation and white matter tract extraction. Voxel time courses were normalized for the purpose of the cmICA computational optimization. Note that temporal variance normalization is usually preferred for connectivity and/or temporal modulation analysis (Elena A. Allen et al., 2012), as it minimizes possible bias in subsequent data reduction.

### 3. Algorithms

#### 1) cmICA

A conventional spatial ICA is typically applied to fMRI data *X*, where we have *Time* by *Space* as input. And ICA separates it into independent spatial sources *S* and their corresponding time-courses *T*, i.e., *X = TS*. Using temporal concatenation of individual datasets, group-level ICA identifies aggregated spatial sources that can be back-projected to estimate subject-specific components, comprised of individual spatial maps *S*_*k*_ and time-courses *T*_*k*_ (V. Calhoun & Adali, 2012). In the case of cmICA, the input connectivity matrix *C* is a *Space* by *Space* matrix, representing the connectivity strength between all node pairs across two brain regions *A* and *B*. And ICA separates it into independent spatial sources *S* in Region *A* and their corresponding connectivity maps *R* in region *B*, i.e., *C = RS* (Wu et al., 2015). Here, *C* can be defined by fiber streamline counts (via tractography) from dMRI between whole brain gray matter as region *A* and whole brain white matter as region *B*, therefore cmICA generates gray matter parcels *S* and white matter tract maps *R* that connected with each *S. C* can also be defined by temporal correlations within whole brain gray matter as both Region *A* and *B*, and cmICA generates gray matter parcels *S* and their functional connected maps *R*.

The use of cmICA to separate a whole brain connectivity matrix is computationally demanding, as it requires an initial computing of voxel-to-voxel correlation (i.e., FC) for the entire brain followed by PCA/ICA on this ultra-large FC adjacency matrix (Fig. 1C). In Wu 2018, we developed an optimization of cmICA to address this problem. Here, we briefly summarize the cmICA algorithm from our previous work (Wu et al., 2015; Wu et al., 2018; Wu et al., 2017a). cmICA was first developed for parcellating tracts in diffusion MRI using a probabilistic tractography connectivity matrix at the voxel scale (Wu et al., 2015). We then extended cmICA to work with fMRI datasets and derived a numerical simplification when the connectivity matrix is defined as the temporal cross-correlation per voxel pair (Wu et al., 2018). This optimization yields spatial sources constrained by FC and their connectivity spatial maps, without actually calculating individual FC matrices.

**Figure 1.**
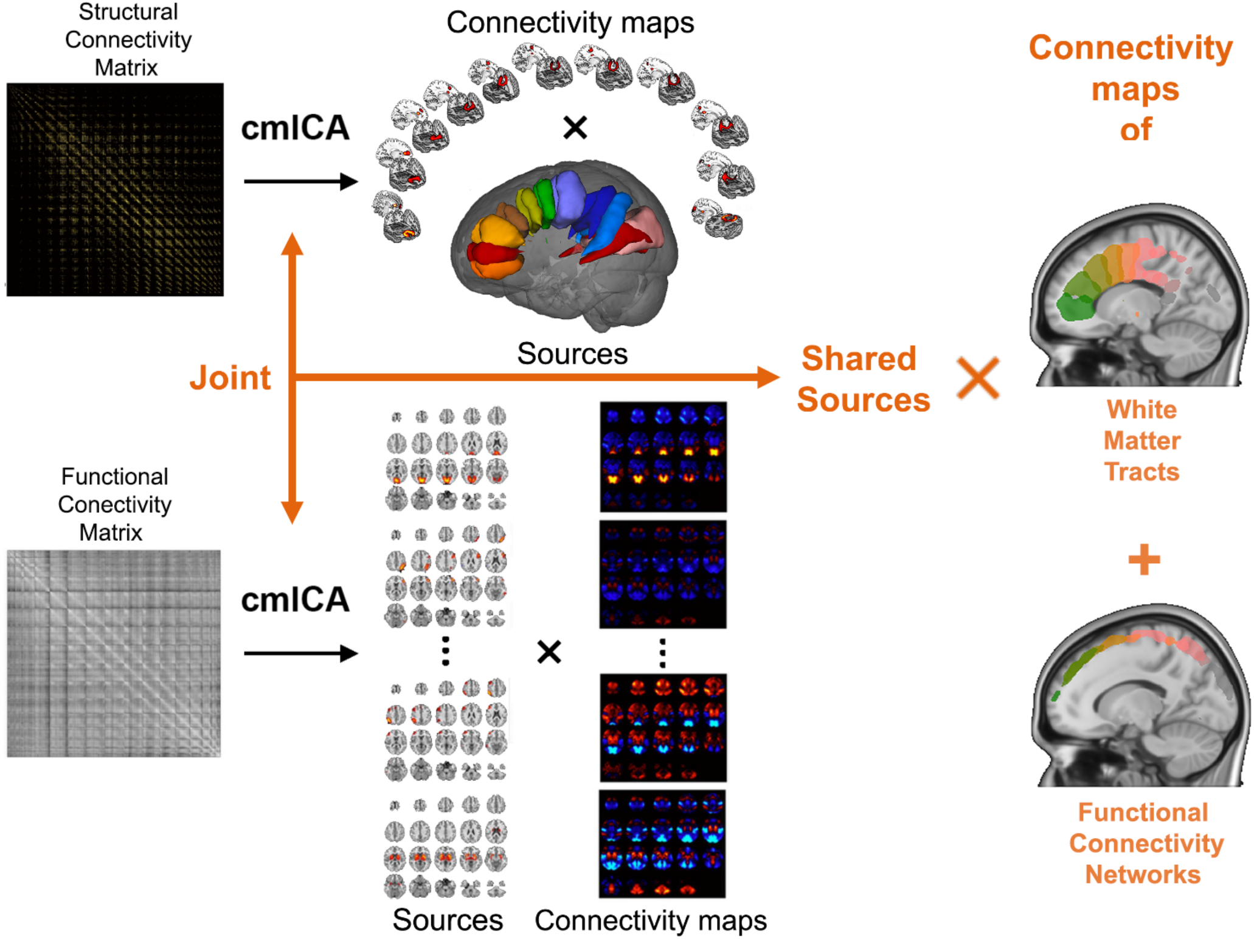
Framework for the proposal joint cmICA approach: cmICA is used to decompose SC matrix and FC matrix into independent sources maps and their corresponding connectivity maps respectively, in top and bottom left. Joint cmICA concatenates two modalities together, extracts shared independent spatial sources from both modalities and their corresponded WM tracts and FC networks.

The advantages of cmICA and its meaning have been discussed in previous work (Wu et al., 2015; Wu et al., 2018). Specifically, we use ICA to decompose each subject’s voxel-wise brain connectivity matrix (*C* = ∑ *S*_*k*_ *R*_*k*_) into a sum of source regions (*S*_*k*_), and their respective connectivity spatial maps (*R*_*k*_) to the whole brain with that source. However, instead of performing a decomposition (source separation) on the original whole brain FC matrix, we simplify it into a straightforward PCA/ICA of the normalized fMRI time series without the need for prior computation on the large FC matrix.

For each single subject, where 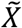 is the ‘temporally normalized’ fMRI for each voxel, we conduct a standard SVD/PCA on 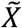. Then,

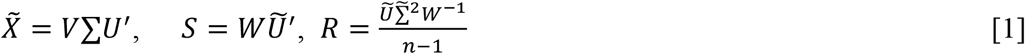

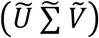 is the reduced SVD/PCA where U and V are the unitary matrices and Σ is the eigenvalue diagonal matrix, W is the ICA demixing matrix.

At the group level, we applied the group ICA approach (Erhardt et al., 2011; Wu et al., 2015)

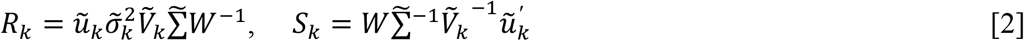

where 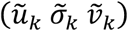 is the reduced SVD/PCA result from single subject *k*, and 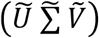 is the reduced SVD/PCA result at concatenated group level where 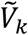 is the partition of 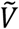 for the subject *k*, and W is the demixing matrix in ICA decomposition.

#### 2) Joint cmICA

In this work we combined two cmICAs, one for temporal-correlation-based functional connectivity (Wu et al., 2018; Wu et al., 2021), and one for tractography-based structural connectivity (Wu et al., 2015), into a single algorithm (Fig. 1). In addition, we enhanced the previous pipeline by adding GPU computation, fast (sparse) PCA and optimization on ultra-large covariance matrices. We used both a FC matrix (gray matter by gray matter as Region *A* by Region *B*) from whole brain fMRI and tractographic SC matrix (gray matter by white matter as Region *A* by Region *B*) from whole brain dMRI as inputs of joint cmICA, by concatenating them together along Region *B* for all subjects. The joint cmICA performs a blind source separation of gray matter independent sources shared between both modalities. Two runs of PCA are conducted, with the first for dimension reduction on the subject-level matrices for both FC matrix and SC matrix and the second for data reduction at group level for two modalities separately prior to performing ICA separation. This latter data reduction balances the differences in the FC and SC data distributions, avoiding ICA selecting ‘sources’ from one modality only due to the significant differences in variance between FC matrix and SC matrix. Finally, joint cmICA generates connectivity-based cortical sources/parcels which are shared between FC matrix and SC matrix. Their corresponding connectivity maps of functionally connected regions (i.e., functional connectivity networks, FC networks) as well as the white matter tracts (WM tracts) that connected these regions, are jointly identified in one estimation using GICA back-reconstruction and concatenation order.

### 4. Model order selection

In this study, we chose a relatively high model order for joint cmICA to effectively capture the information from both modalities, as well as to match the model dimensions that we have validated in our previous work (Abou-Elseoud et al., 2010; Kiviniemi et al., 2009; Wu, Eichele, & Calhoun, 2010). In Wu 2010, we found 30 reliable cmICA sources of white matter tracts in COBRE data. We also chose to use a symmetrical PCA/ICA model from two modalities, therefore, a model order of 60 was selected.

To ensure the validity of model order selection on our dataset, ICASSO (Himberg, Hyvärinen, & Esposito, 2004) with 20 re-runs and random initial conditions was used to test the convergence during ICA training and the stability and the ‘best run’ was selected to provide a robust and replicable result using a minimum spanning tree (MST) method introduced in (Du, Ma, Fu, Calhoun, & Adali, 2014) as implemented in the GIFT toolbox (http://trendscenter.org/software/gift).

### 5. Feature Selection

Joint cmICA resulted in 60 independent source maps which were further processed and visualized using the following procedure (Elena A Allen et al., 2011; V. D. Calhoun, Wu, Kiehl, Eichele, & Pearlson, 2010; Wu et al., 2018; Wu et al., 2010). First, the components were sorted by mean variances, i.e. its ‘significance’/’contribution’ towards input data. Second, as a large variance imbalance between FC matrix and SC matrix was observed, each modality’s contribution towards final cmICA sources would be different. To ensure shared sources from both modalities were properly selected, a contribution ratio was computed by averaging the absolute value of loadings of each component for each modality separately, and then calculating the proportion from one to another. In this study, no more than 80% and no less than 20% contributions were considered meaningful ‘shared’ component. We’ve also examined the spatial correlations between the source maps from two modalities for both group average and individual maps, and ensured the significance of these correlations. A routine motion, imaging and physiological artifact check was also performed (Elena A Allen et al., 2011; Mantini, Corbetta, Perrucci, Romani, & Del Gratta, 2009).

Furthermore, we also compared the sources generated from the joint cmICA model and sources generated from unimodal cmICA estimated separately on fMRI and dMRI, to examine whether cortical parcellations are different from the two methods.

## Results

### 1. Matching gray matter parcellations from FC matrix and SC matrix

Joint cmICA yields aggregated gray matter S sources along with FC matrix and SC matrix’s ‘contribution’ towards the joint sources for evaluations. After back-reconstruction via Equation [2], group-averaged gray matter sources from each modality are used for comparison and evaluation.

After removing artifactual components as well as components that were not shared between two modalities from feature selection (see methods), 28 sources were chosen for further analysis. Here we show the 28 selected dual *S* maps from FC matrix and SC matrix in Figure 2 and in MNI details in Table 1. The remaining components are listed in the supplemental. We grouped sources into 5 sub-categories based on the linked white matter tracts of *R* maps from SC matrix (Figure 4-6, (Wu et al., 2015). Figure 2 (A-E) showed the gray matter sources map connected by WM tracts of commissural fibers (CF), inferior longitudinal fasciculus (ILF) / inferior frontal-occipital fasciculus (IFOF) / Uncinate fasciculus (UF), superior longitudinal fasciculus (SLF), cingulate (CG), and corticospinal tracts (CST), respectively (see details in Results 2). Each *S* map was thresholded for visualization purposes by a suitable threshold. Component t-statistic maps were computed and thresholded at *t* > μ + 3σ, with μ being the mean and σ the standard deviation of spatial component (as discussed by (Elena A Allen et al., 2011)).

**Table 1.**
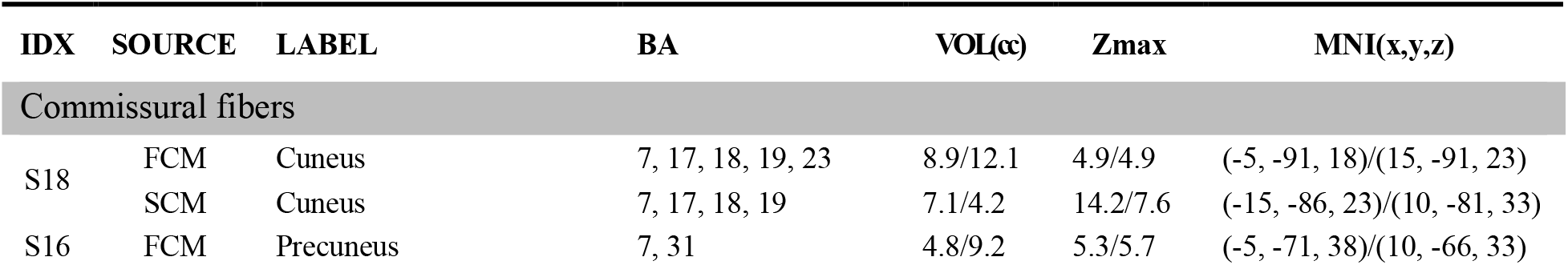
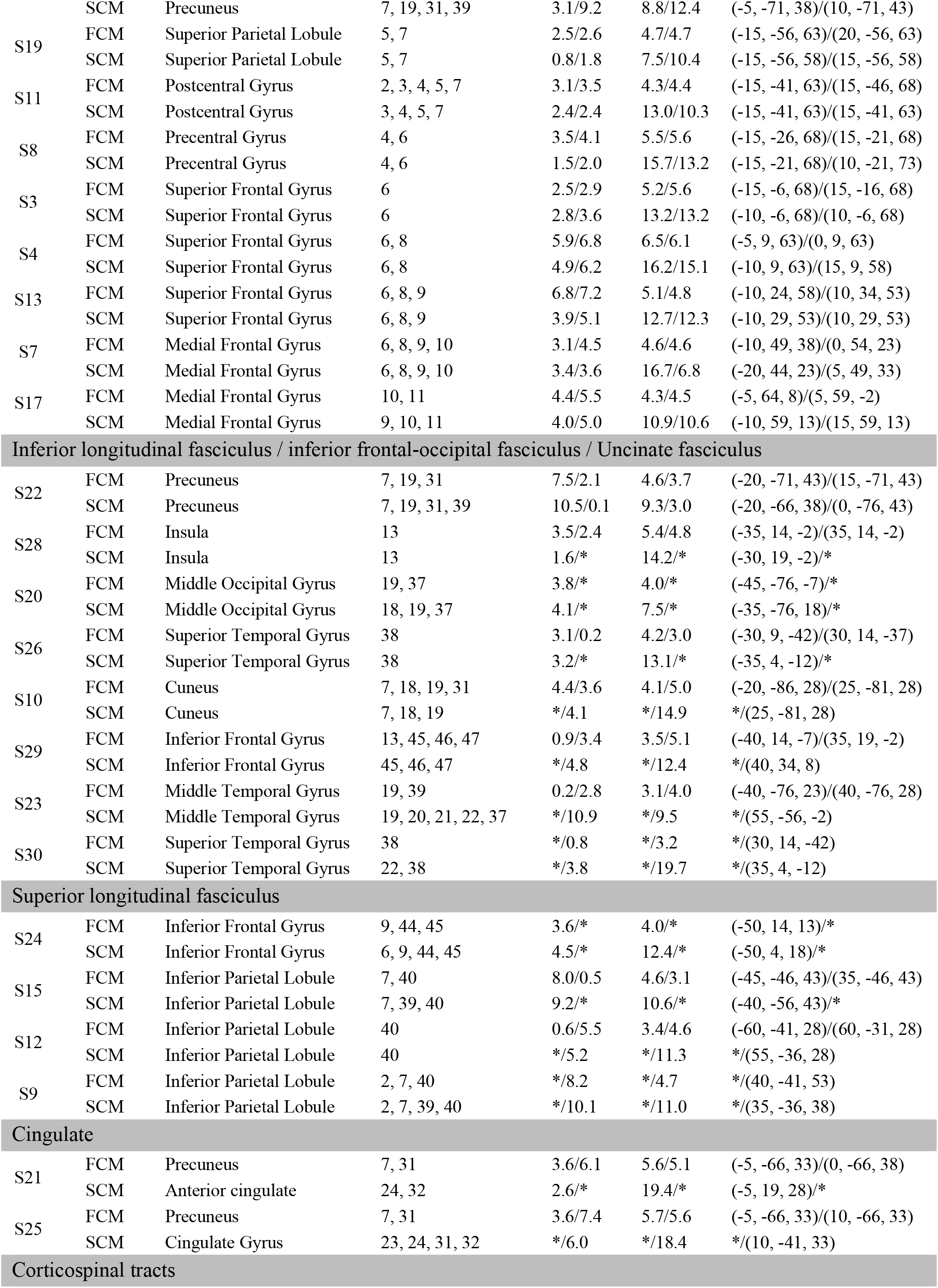
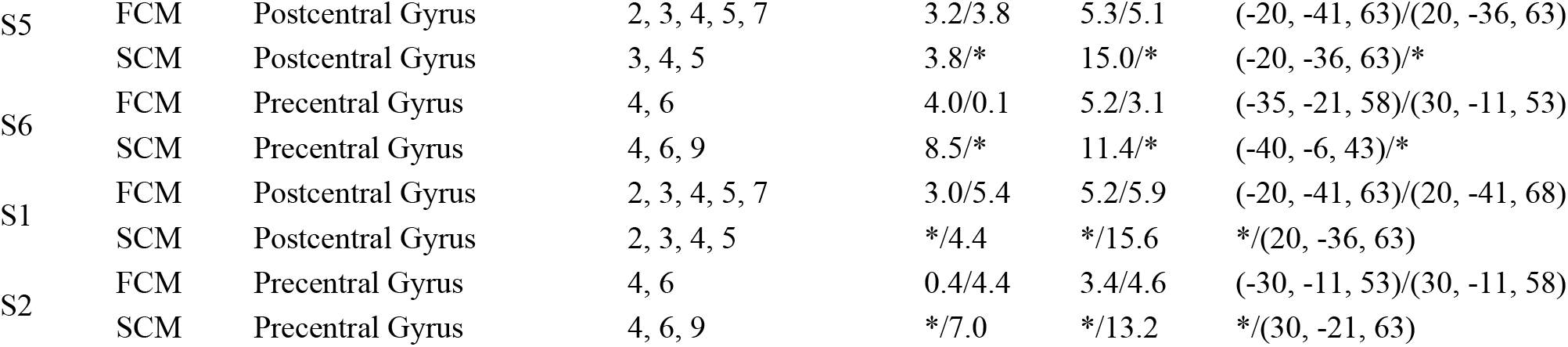
MNI table of joint cmICA sources *S*

**Figure 2.**
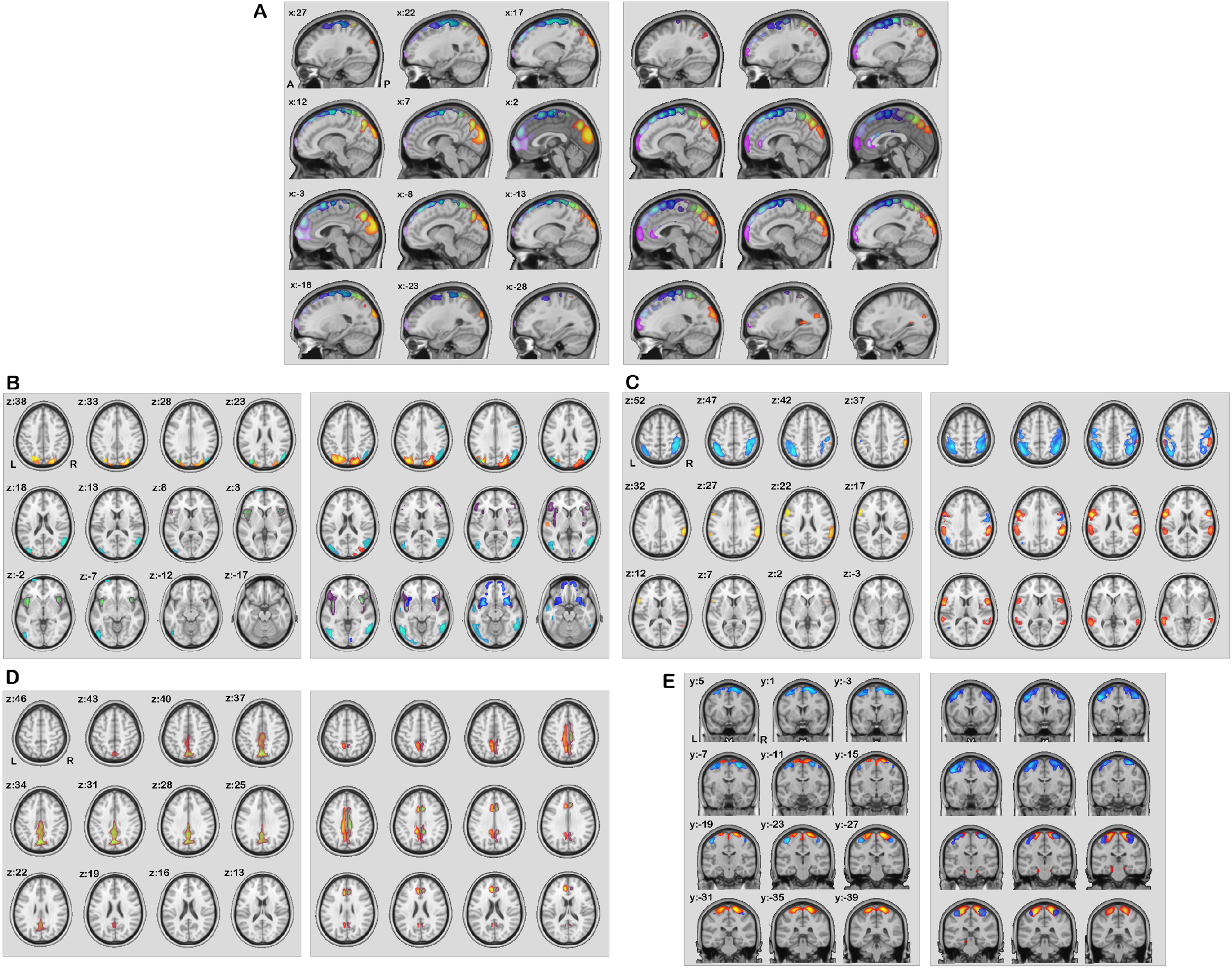
Gray matter sources from FC matrix and SC matrix: All sources *S* are grouped by their connected WM tract *R*’s category (indicated in Figure 4-6). (A-E) show the five types of cortical sources connected by commissural fibers (CF), inferior longitudinal fasciculus (ILF) / inferior frontal-occipital fasciculus (IFOF) / Uncinate fasciculus (UF), superior longitudinal fasciculus (SLF), cingulate (CG), and corticospinal tracts (CST), respectively. In each sub-plot, left shows the sources back-reconstructed from FC matrix and right shows the ones from SC matrix.

Although connectivity input from two different modalities were balanced out by selecting equal share of PCA results prior to ICA source separation, we found that the final outcome is not balanced, i.e., each ICA sources are biased towards one modality or another. Results showed that, after sorting by variance and contribution ratio between both FC matrix and SC matrix, the first half of 30 sources were contributed more from SC matrix, whereas the second half of 30 sources were more from FC matrix (see supplemental). This dichotomy here is caused by spatial distribution differences from two very different modalities that are not influenced by balancing variance, rescaling etc. More importantly, the first 30 sources were much more evenly contributed between two modalities than the second half (Figure 3A and Supplemental). And in this study, we focused on more ‘shared’ sources across two modalities, which passed the contribution criteria (Figure 3A). In addition, spatial similarities between each source pairs are shown in Figure 3B.

**Figure 3.**
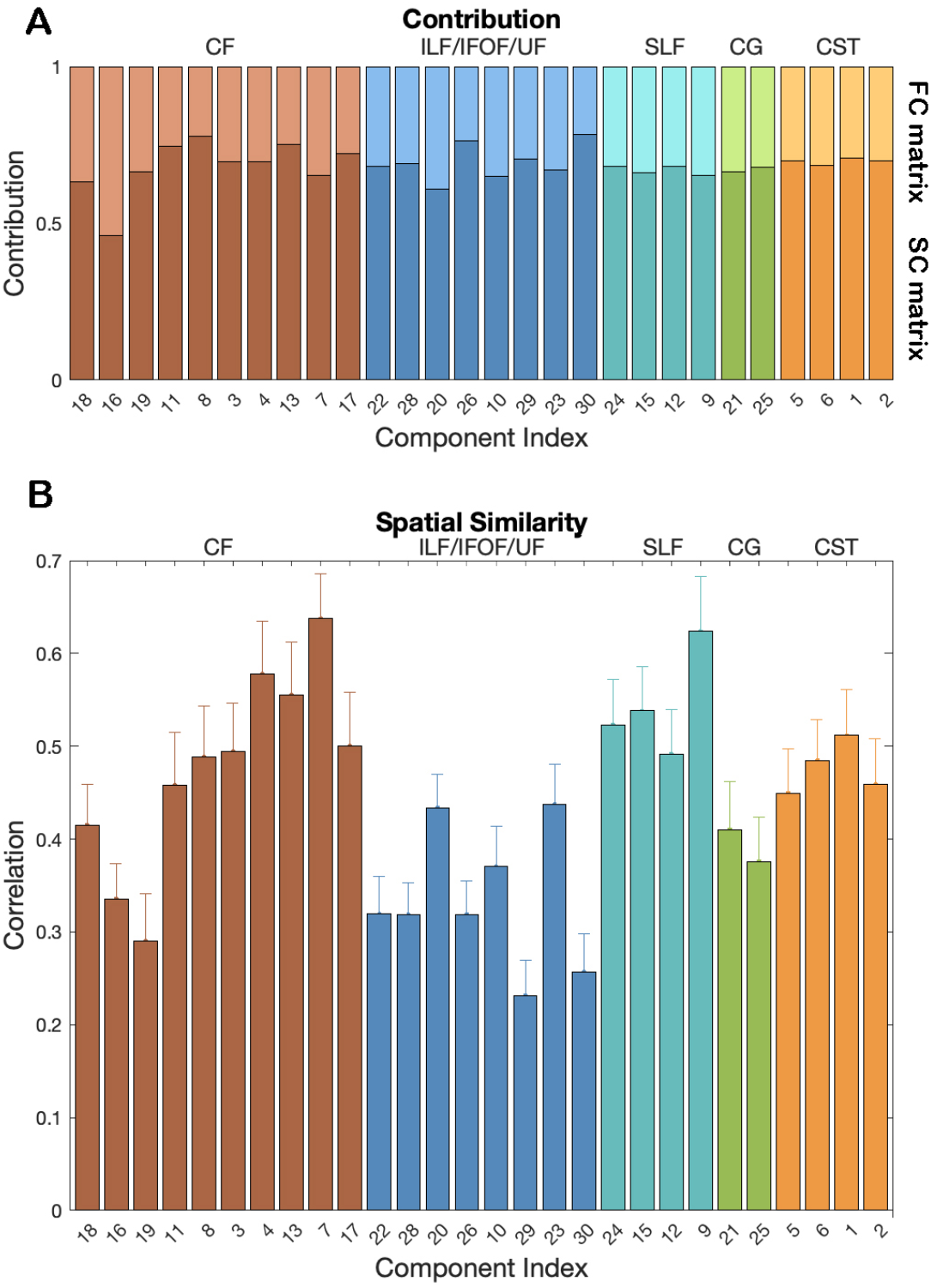
Contribution ratio and spatial similarity of source maps between two modalities: (A) shows the 30 most shared sources and the contribution ratio from two modalities. (B) shows the spatial similarities (correlations) between each source pairs.

### 2. Match functional connectivity networks and white matter tracts

As mentioned above, we found the components more shared between SC matrix and FC matrix are slightly biased towards SC matrix’s contribution. Furthermore, we found that the resulting 28 components’ connectivity *R* map from SC matrix were well matched with white matter tracts from our previous work (Wu et al., 2015), we followed the same grouping of 28 spatial maps into three major functional categories as before, consisting of commissural (right-left hemispheric cortex), association (same hemispheric cortex-cortex) and projection (cortex-spinal, cortex-thalamic) fibers (Mori, Wakana, Van Zijl, & Nagae-Poetscher, 2005; Wakana, Jiang, Nagae-Poetscher, van Zijl, & Mori, 2004), with names in each category determined by the 20 region ‘JHU white-matter tractography atlas’ (FSL-Atlases; Hua et al., 2008; Wakana et al., 2007). Here, we jointly showed connectivity maps *R* from FC matrix i.e., FC networks, and *R* from SC matrix i.e., WM tracts, derived from shared gray matter sources *S*. Note that, the *R* maps for FC networks are not the same maps as the *S* maps for either modality. These maps do not have the restriction of independence or stationarity as the *S* maps, and mathematically they can be seen as a seed-based (correlation) connectivity maps of *S* as a ROI (Wu et al., 2018). The cmICA connectivity maps *R* were thresholded for visualization using *t* > μ + 2σ, due to a less sharp spatial distribution than *S* (Wu et al., 2018). Individual component details are listed in the supplemental.

In Figure 4, the commissural fibers (darker colored in multi-component composite) define the fibers that go across the corpus callosum connecting both hemispheres of the brain. We found that 10 out of the 28 ICA components belonged to this category. We also found that each oriented commissural component connects different cortical areas (translucent matching colored in multi-component composites) across the two hemispheres.

**Figure 4.**
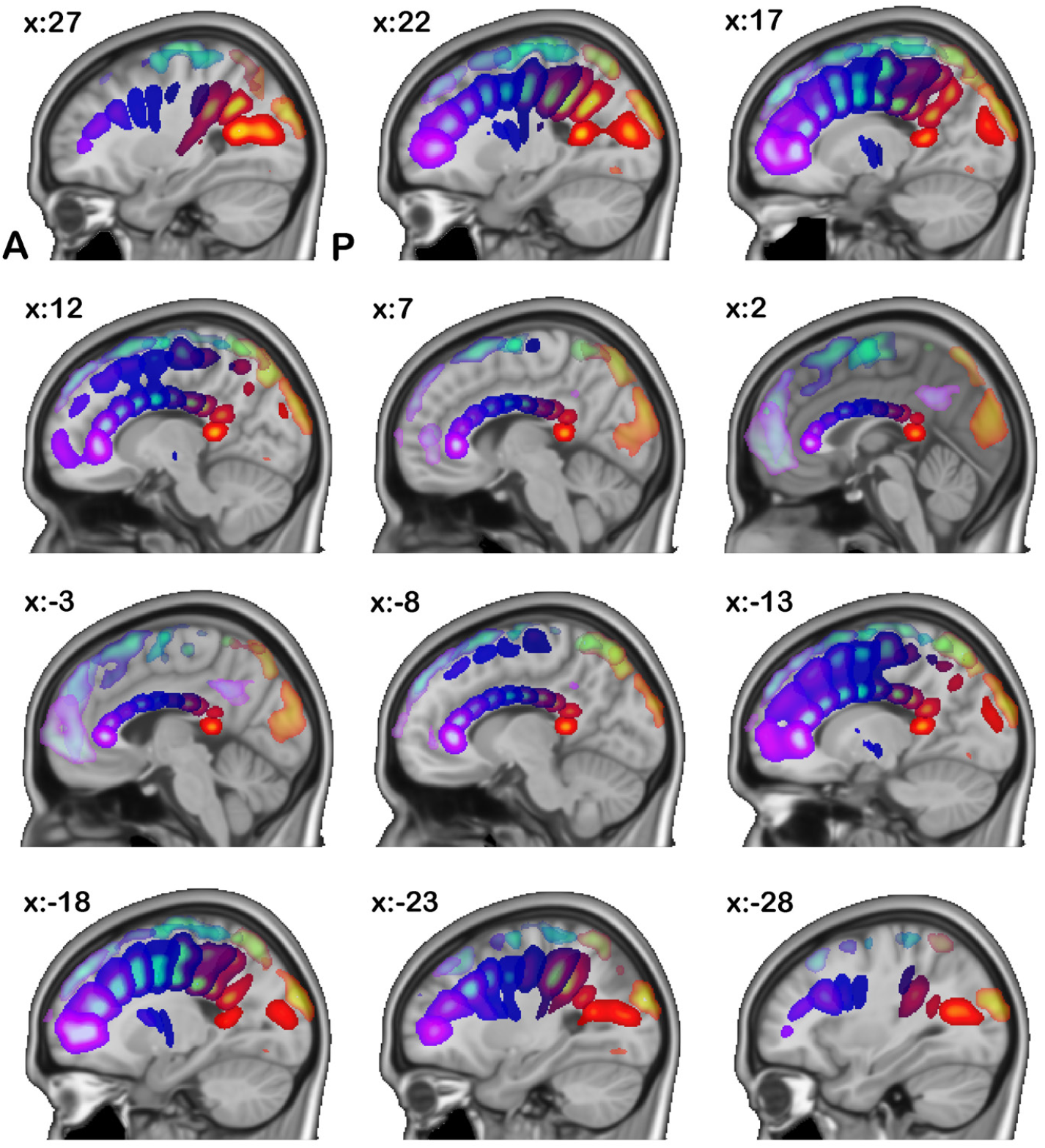
: Commissural components: Here we show components grouped into the commissural fiber category based on their numerical correspondence to the JHU white matter atlas. We show both the FC network *R* maps (translucent lighter colored) from fMRI and their corresponding WM tract *R* maps from dMRI (deeper matching colored). Each pair of left/right components are showed in the same color.

Joint cmICA segmented the corpus callosum into 10 segments based on the connectivity of these regions from SC matrix, a well-behaved fanning pattern, which is consisted with our previous findings (Wu et al., 2015) and Hofer’s tractography based findings (Hofer & Frahm, 2006) but with a finer corpus callosum segmentation. More importantly, results also identified the corresponding cortical FC networks that were linked to these white matter commissural tracts.

The association fibers connect different parts of the brain within the same cerebral hemisphere. These include Figure 5A the inferior longitudinal fasciculus (ILF), inferior frontal-occipital fasciculus (IFOF) and uncinate fasciculus (UF), Figure 5B superior longitudinal fasciculus (SLF, temporal and parietal), and Figure 5C cingulum (superior cingulate, supracallosal). All these tracts connect different part of the cortical areas longitudinally within the same hemisphere instead of across two hemispheres. Left/right (lateral) component pairs are plotted in the same color.

**Figure 5:**
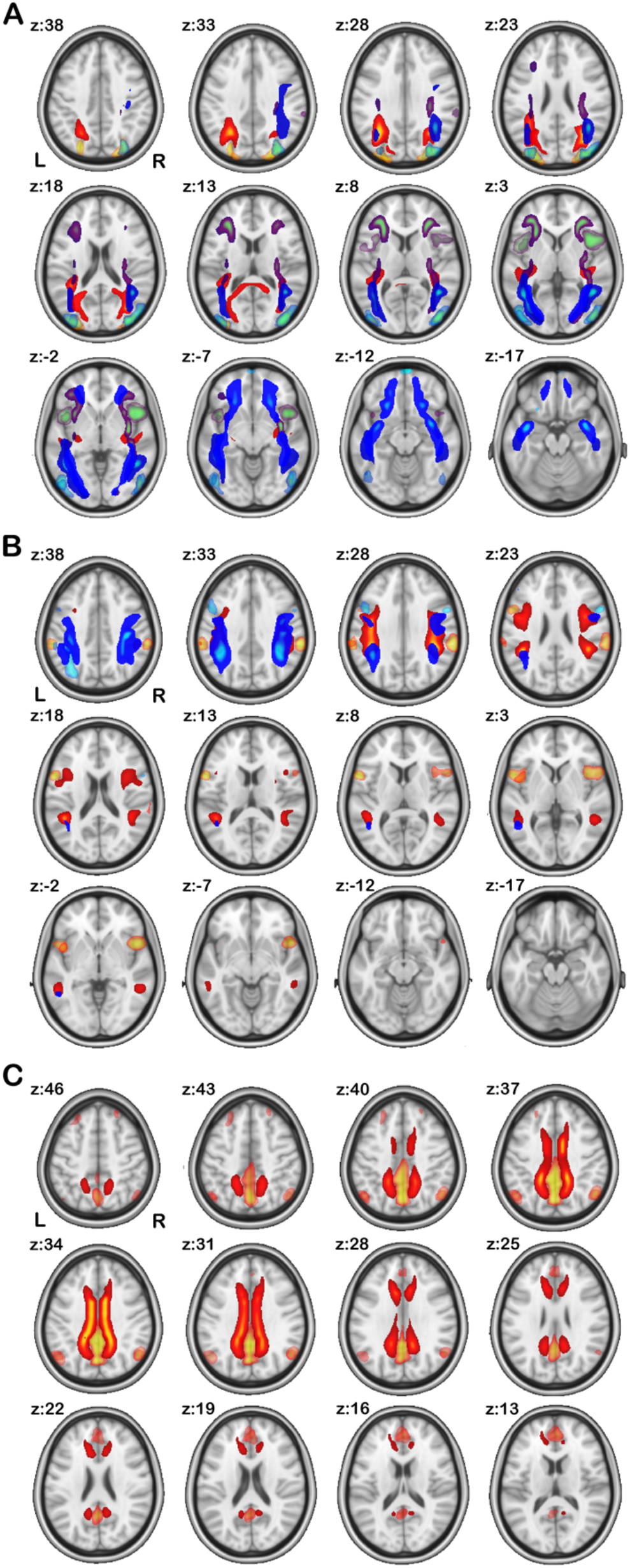
Association components: Here we show components grouped into the association fiber category based on their numerical correspondence to the JHU white matter atlas. We show both the FC network *R* maps (translucent lighter colored) from fMRI and their corresponding WM tract *R* maps from dMRI (deeper matching colored). Each pair of left/right components are showed in the same color.

In Figure 5B parietal cortex and frontal/precentral cortex are connected by SLF in both hemispheres. Also note that in Figure 5C both left and right cingulate tracts (bright hot colored) connect the long-range ACC and PCC (translucent hot colored) that are linked to the conventional default mode network (DMN). The angular gyri in DMN are also shown in FC network in Figure 5C, but no connection is shown in WM tracts from angular gyrus towards the other DMN areas. It is possibly because 1) PCA/ICA only extracts the ‘major’ white matter tract bundles, i.e., high fiber counts, whereas the white matter fibers that connects angular gyrus (in DMN) have much ‘weaker’ connectivity/fiber counts 2) the extracted WM tract and FC network *R* maps are guided by source map *S*, not from each other. Comparing Figure 5C with Figure 2D, it is noticed that S 21 and S 25 are located mostly in anterior/posterior cingulate from both FC and SC with no coverage of angular gyrus, therefore, R21 and R25 from SC show a cingulum only because it is the most structurally-connected (fiber counts) tracts to these areas, but R 21 and R 25 from FC are not restricted in just cingulate but also covering angular gyri, because the FC network is based on temporal correlation and angular gyri are highly functionally connected (correlation) with S 21 and S 25 too.

The projection fibers connect the cortex to the lower parts of the brain and spinal cord, and include the corticospinal tracts. In Figure 6, corticospinal tracts originate in primary motor cortex and somatosensory cortex and descend into brainstem, serving as the major pathways for carrying movement-related information. Following Figure 5, each left/right pair of components are plotted in the same color.

**Figure 6:**
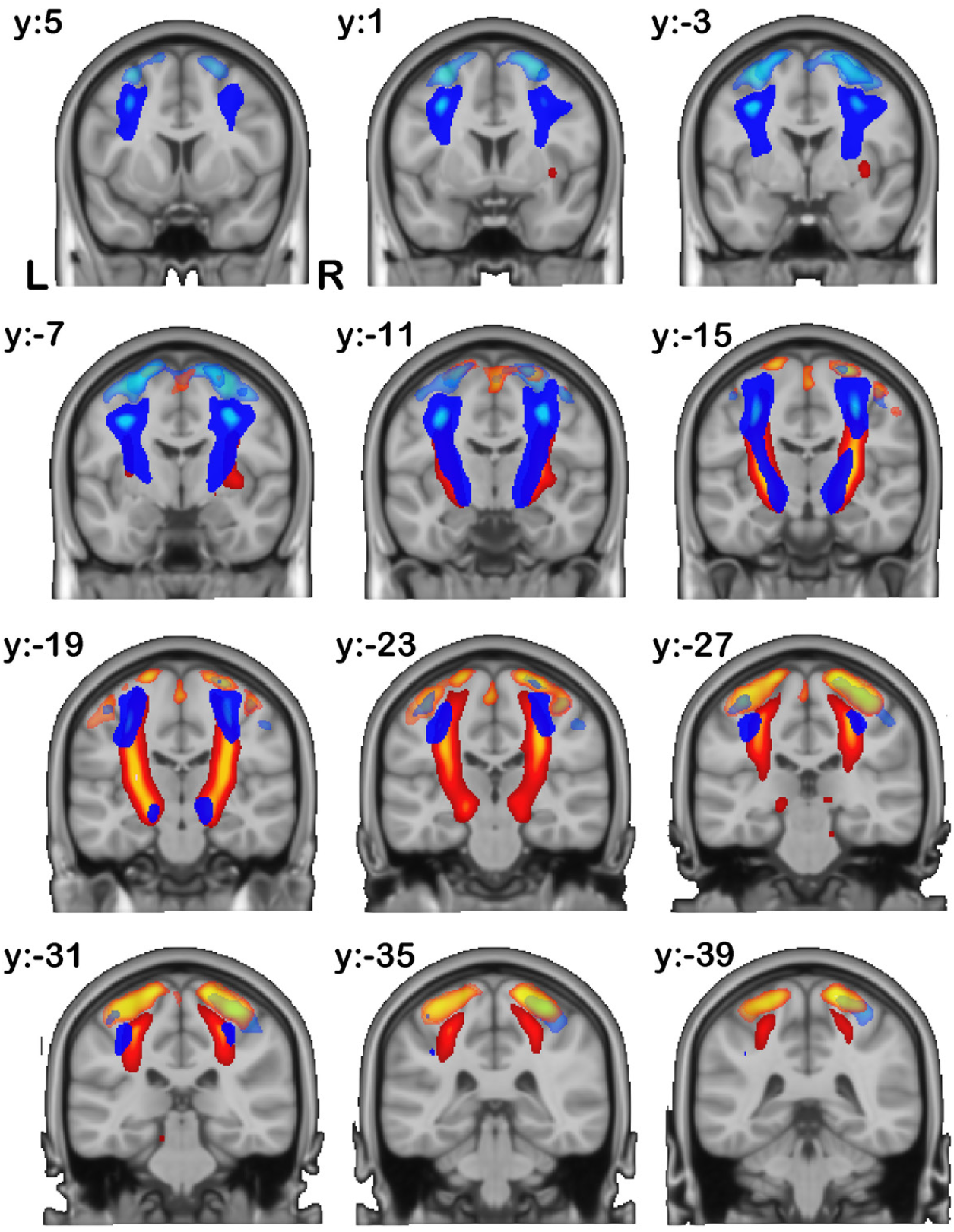
Projection components: Here we show components grouped into the projection fiber category based on their numerical correspondence to the JHU white matter atlas. We show both the FC network *R* maps (translucent lighter colored) in gray matter from fMRI and their corresponding WM tract *R* maps in white matter from dMRI (deeper matching colored). Each pair of left/right components are showed in the same color.

The spatial tract components were numerically compared to the JHU 20 region white matter tract atlas, based on the overlap of the ICA components to the JHU atlas regions. Table 2 shows the percentage overlap of ICA maps converted to an ROI to the JHU atlas (*JHU-ICBM-tracts-maxprob-thr0-1mm* from the FSL atlas library). Note that, JHU atlas is only used as a reference to name tract bundles in Table 2, and we labeled all the fiber bundles according to the atlas region with which they had maximum correspondence, regardless the actual fiber orientations. JHU 20 region tract atlas does not include all fanning-out commissural components, instead it only has ‘Forceps major’ and ‘Forceps minor’. In the cases of R13, it is in fact a commissural tract based on its orientation and shape, instead of ‘Cingulum’ which is an association tract. Yet it matches with Cingulum L in JHU tract Atlas with a very low (11.3%) overlap, because it is still the highest match compared to the other 19 JHU regions. The same applies to R16, R19, R11, R8, R3, R4, with very low overlaps with various atlas regions, as JHU atlas does not cover these commissural fibers as well. The ‘true’ Cingulum pairs are R21 and R25, and ‘true’ SLFs are R24, R15, R12 and R9. This results also highlights the benefits of using a data driven approach, as existing atlases do not always well capture the complexity of existing tracts or functional networks.

**Table 2.** White matter tract *R* maps’ correspondence to JHU tractography atlas

| Index | Fiber type | JHU Atlas Region | % |
| --- | --- | --- | --- |
| R18 | Commissural | Forceps major | 85.3 |
| R16 | Commissural | Inferior fronto-occipital fasciculus R | 16.7 |
| R19 | Commissural | Anterior thalamic radiation L | 12.0 |
| R11 | Commissural | Corticospinal tract L | 15.9 |
| R8 | Commissural | Corticospinal tract R | 19.0 |
| R3 | Commissural | Superior longitudinal fasciculus L | 5.7 |
| R4 | Commissural | Superior longitudinal fasciculus L | 5.4 |
| R13 | Commissural | Cingulum (cingulate gyrus) L | 11.3 |
| R7 | Commissural | Forceps minor | 61.0 |
| R17 | Commissural | Forceps minor | 75.4 |
| R22 | Association | Inferior fronto-occipital fasciculus L | 31.0 |
| R28 | Association | Inferior fronto-occipital fasciculus L | 60.0 |
| R20 | Association | Inferior longitudinal fasciculus L | 51.0 |
| R26 | Association | Inferior fronto-occipital fasciculus L | 33.5 |
| R10 | Association | Inferior fronto-occipital fasciculus R | 32.8 |
| R29 | Association | Inferior fronto-occipital fasciculus R | 65.0 |
| R23 | Association | Inferior fronto-occipital fasciculus R | 43.9 |
| R30 | Association | Inferior fronto-occipital fasciculus R | 41.6 |
| R24 | Association | Superior longitudinal fasciculus L | 94.1 |
| R15 | Association | Superior longitudinal fasciculus L | 84.8 |
| R12 | Association | Superior longitudinal fasciculus R | 89.8 |
| R9 | Association | Superior longitudinal fasciculus R | 89.8 |
| R21 | Association | Cingulum (cingulate gyrus) L | 79.5 |
| R25 | Association | Cingulum (cingulate gyrus) R | 68.7 |
| R5 | Projection | Corticospinal tract L | 59.3 |
| R6 | Projection | Superior longitudinal fasciculus L | 43.2 |
| R1 | Projection | Corticospinal tract R | 62.9 |
| R2 | Projection | Superior longitudinal fasciculus R | 38.0 |

## Discussion

The proposed joint cmICA approach extracts gray matter sources from both SC and FC information, and generates both the functional network connectivity map and white matter tract connectivity map, all in one scheme. To our knowledge, this is the first approach that jointly extracts connectivity-based sources from whole-brain fMRI and dMRI without providing any prior information.

### 1. Linking structural connectivity and functional connectivity

In prior studies, the most widely used approaches to match SC and FC involves evaluating the correlation/distance between (vectorized) SC and FC matrices (Honey et al., 2009; Liégeois, Santos, Matta, Van De Ville, & Sayed, 2020; Suárez-Méndez et al., 2020), where both matrices’ nodes are pre-defined and macroscopic.

Using cmICA, however, eliminates the need for prior selection of ROIs for both modalities, which may be biased and sensitive to noise especially in group analysis (Van Den Heuvel & Pol, 2010). ICA has achieved great success in neuroimaging application, particular in fMRI (V. Calhoun & Adali, 2012; V. D. Calhoun & Adali, 2006; McKeown, Hansen, & Sejnowsk, 2003). ICA is typically used to extract sources that are purely data-driven by maximizing spatial independence. In our case, we sought cortical sources that were both spatially independent and best shared between SC and FC, meanwhile recovering these sources’ conjugate connectivity profile maps in FC and SC, yielding FC networks and WM tracts respectively. This approach opens avenues to explore functional connectivity and structural connectivity maps in a data-driven way, which we believe is useful and straightforward, compared to other parcellation methods requiring details in clustering priors and/or projections (Honey et al., 2009; Iraji et al., 2016; O’Muircheartaigh et al., 2011).

### 2. ICA based joint analysis

Using ICA to combine fMRI and dMRI analysis is not new. Other approaches such as joint-ICA, parallel-ICA, or more extended ICA/canonical correlation analysis (CCA), or independent vector analysis (IVA) have been used to seek connections between two modalities (Sui, Adali, Li, Yang, & Calhoun, 2010). However, this is different from our proposed method, as previous ICA related methods do not involve ‘connectivity’ directly. These methods used features, e.g., fractional amplitude of low frequency fluctuation (fALFF) maps or regressed activation maps from fMRI, and fractional anisotropy (FA) maps or other tensor metrices from dMRI, as input of multi-model analysis, and extracted shared information along subjects’ loadings (V. D. Calhoun & Adali, 2008; V. D. Calhoun & Allen, 2013). This imposes two potential problems. First, using different features may yield very different links among the multimodal sources. Second, in many cases, the linked sources from two modalities are not anatomically aligned and/or connected. Previous approaches do not link anatomic and functional sources automatically, making the interpretation and validation of the results more difficult. Our approach, in contrast, automatically identifies the shared cortical sources from FC and SC, and the results of their connectivity maps in gray matter and white matter, without spatial constraints in computation.

### 3. Multimodal cortical parcellations

Cortical organization and mapping has been an essential objective in neuroscience for over a century, where it serves as models for human brain function in development, aging, health and disease (Glasser et al., 2015; Van Essen et al., 2019). Such topic has been most commonly analyzed using surface-based approaches to reveal the topology of the cortical sheets (Thyreau & Taki, 2020; Van Essen et al., 2019). However, accurate delineation of the entire mosaic of cortical areas requires extensive modeling and assumptions, whereas newer methods, such as deep learning network, replies on tremendous training on large data scale for reliability (Fedorov et al., 2017; Thyreau & Taki, 2020). In contrary, our approach makes minimal assumptions and gives an instant visual perspective of cortical organization as well as their axonal linkages.

Prior work has also indicated that a multimodal but integrated approach that utilizes information from anatomy, structure, function, connectivity and topological organization would work better in terms of cortical connectivity analysis, as each modality provides complementary information and benefits (Glasser et al., 2015; Van Essen et al., 2019). In this work, our results clearly showed that different cortical parcels, which spread from prefrontal, premotor, motor and sensory, to parietal, temporal occipital regions, were connected by differently oriented commissural fiber bundles across corpus callosum. And the default model network, typically only recovered from functional connectivity analysis, was also detected in tract-based connectivity linked by a pair of longitudinal cingulate tracts.

### 4. Limitations and future work

This study is primarily exploratory. Our results showed that ICA cortical sources derived from both SC and FC data are not symmetrically contributed from different modalities. Cortical sources that are shared more often between two modalities are more often contributed by SC, whereas cortical sources with more FC contributions are shared less often. Comparing our two separate cmICA works in SC and FC (Wu et al., 2015; Wu et al., 2018) and knowing that ICA decomposition is driven by spatial distribution, it suggests the reason behind is that spatial distribution from two different modalities are quite different, specifically, the SC matrix is defined by fiber streamline counts and the FC matrix is defined by temporal correlations. The former has a much more sparse distribution and more stable sources than the latter. This suggests that the spatial distribution differences between these two modalities are linked to the different ways in which the sources decompose and how they share from each other. However, it is important to note that joint cmICA itself is not SC guided or a biased modeling method. Rather, joint cmICA treats the contributions from each part of the modalities equally. If the data from two modalities have no radical differences in their spatial distributions, joint cmICA will result in a balanced source separation.

Although we were able to address certain degree of contribution bias by balancing PCA outputs from both modalities, the current model has not created an equally contributed ICA decomposition from both SC and FC due to their drastic spatial distribution differences. Yet, upon testing, enforcing the loading/contribution to be identical (e.g., in joint-ICA) or as high as possible (e.g., searching a gradient descendent to seek highest correlation between two modalities) will impact the independence/sparsity between sources in spatial distributions and reduce the quality of the white matter tract results since the latter is along the loading direction. To further balance out the intrinsic data differences and maintain component quality, a more sophisticated fusion strategy, using either a constrained decomposition or possibly leveraging non-linear relationships (Motlaghian et al., 2021), can be leveraged and will be explored in our future work. There is also some existing model-based work between SC and FC, e.g., (Chu et al., 2018), which has the potential to be extended into joint cmICA.

The matching of sources from SC and FC as well as FC network and WM tract at the individual subject level was not the focus on this study. However, it is worth investigating the specific correspondence and differences in subjects. In our prior work (Wu et al., 2015; Wu et al., 2018; Wu et al., 2021), we showed that cmICA is sensitive to individual variability, which suggests individual subject joint matching could be done using dual model ‘fitting’ at the subject level.

Last, but not least, a challenge in implementing joint cmICA and cmICA on SC is the large computation and storage cost of tractography at whole brain to whole brain voxel scale. For large studies this requires a substantial amount of CPU/GPU resources to run within a reasonable time frame.

## Conclusion

In this study, we proposed a data-driven joint cmICA to integrate and analyze structural connectivity and functional connectivity simultaneously, systematically, and conveniently, without the need for a prior atlas. The decomposed connectivity parcels indicate a shared cortical segregation from both structural organization and functional connectivity, though with various degrees of (dis)similarity. Moreover, we show conjoint structural WM tracts and FC networks that directly link these cortical parcellations/sources, and explain to some degree how the structural fiber firing connects the functional networks.

Joint cmICA incorporates both structural and functional connectivity, which can be complementary to each other and may detect disease traces that could be missing by using either modality alone. In addition, as connectivity maps R’s are extracted from the loading parameters rather than the sources, they are not restricted by an assumption of independence or stationarity. This may significantly increase the sensitivity to capture individual variability in both healthy subjects and in diseases, e.g. schizophrenia, where group differences between controls and patients are very subtle, and might not be well detected by feature-based ICA models. Overall, joint-cmICA provides an effective tool for connectivity-based multimodal data fusion in brain, and has great potential for application to study both the healthy and disordered brain.

## Acknowledgement

This work was funded by the National Institutes of Health (NIH) grants 1R01EB006841, 1R01EB005846, and the National Science Foundation (NSF) grant #2112455.

## Data Availability Statement

The data that support the findings of this study are available from the corresponding author upon reasonable request or from COINS (http://coins.trendscenter.org) (Scott et al., 2011).

