## supplemental for "Joint cmICA: auto-linking of structural and functional connectivity"

1. 60 source maps' contribution from FC matrix and SC matrix

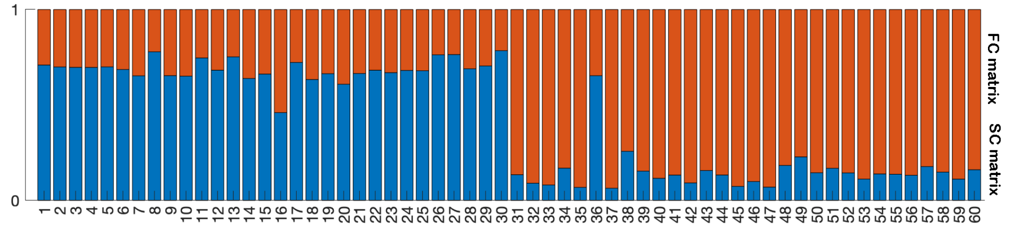

2. Joint cmICA *S* parcellations from FC matrix ('red') and SC matrix ('blue'), all 60 components.

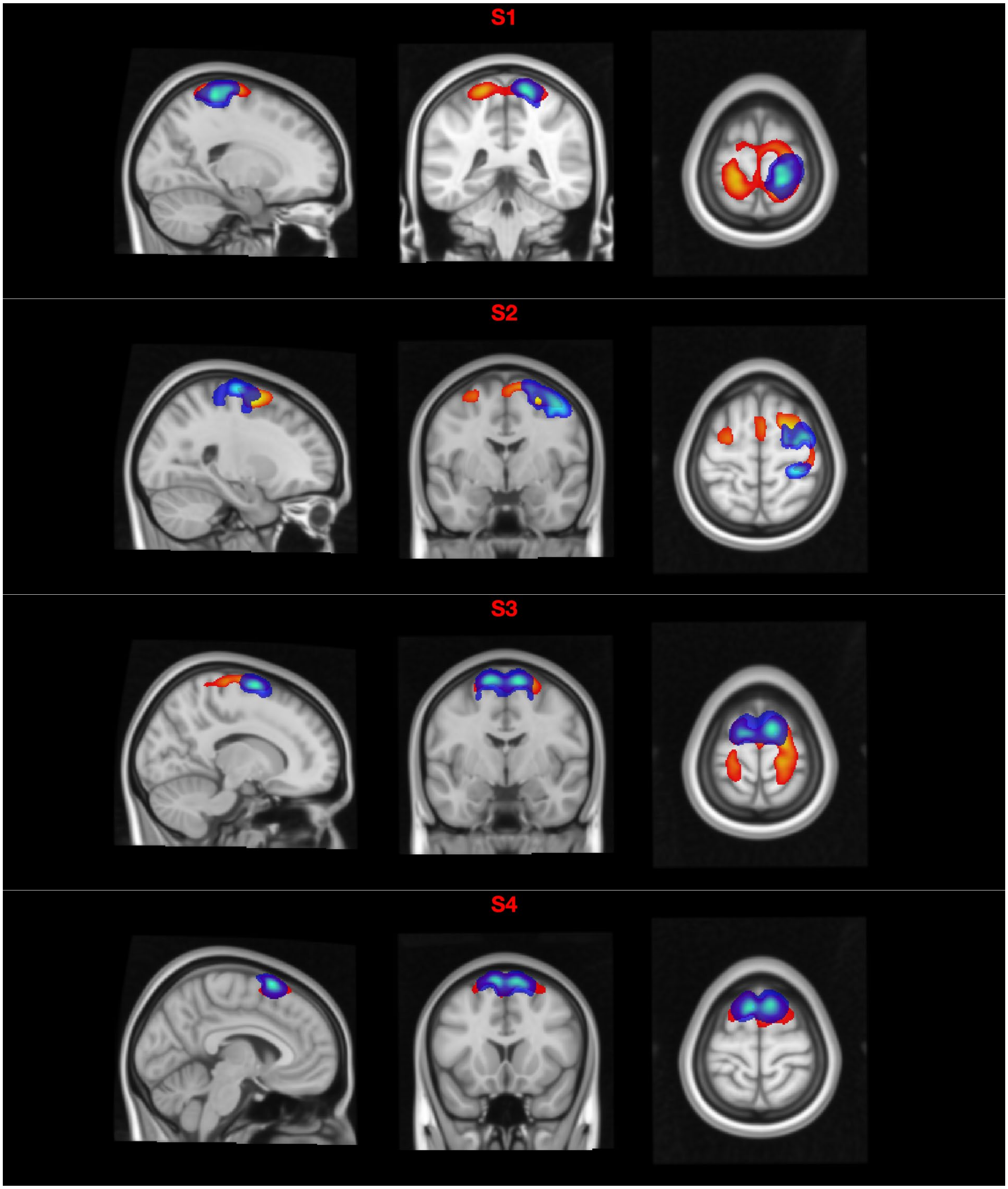

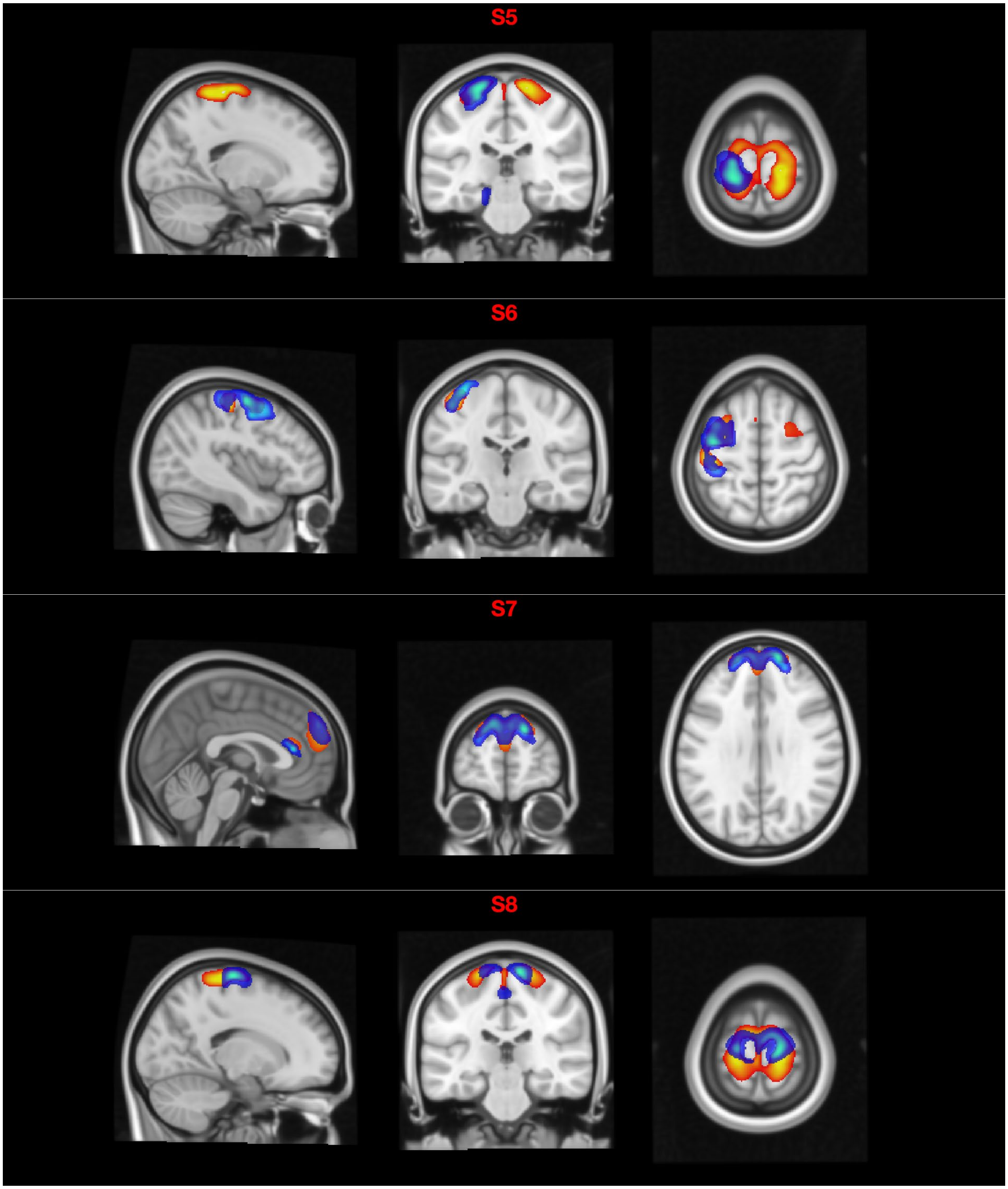
S5
S6
S7
S8

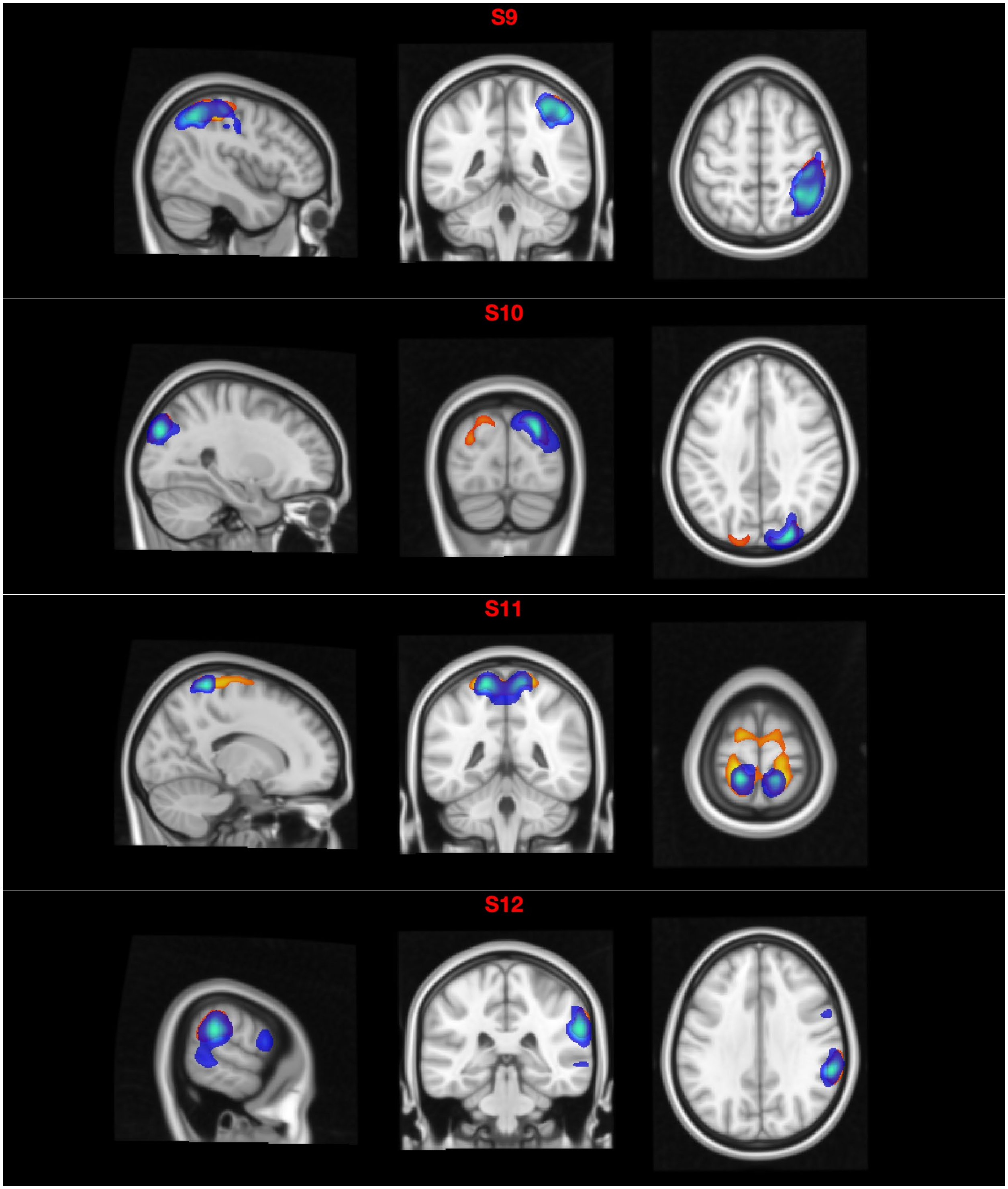
S9
S10
S11
S12

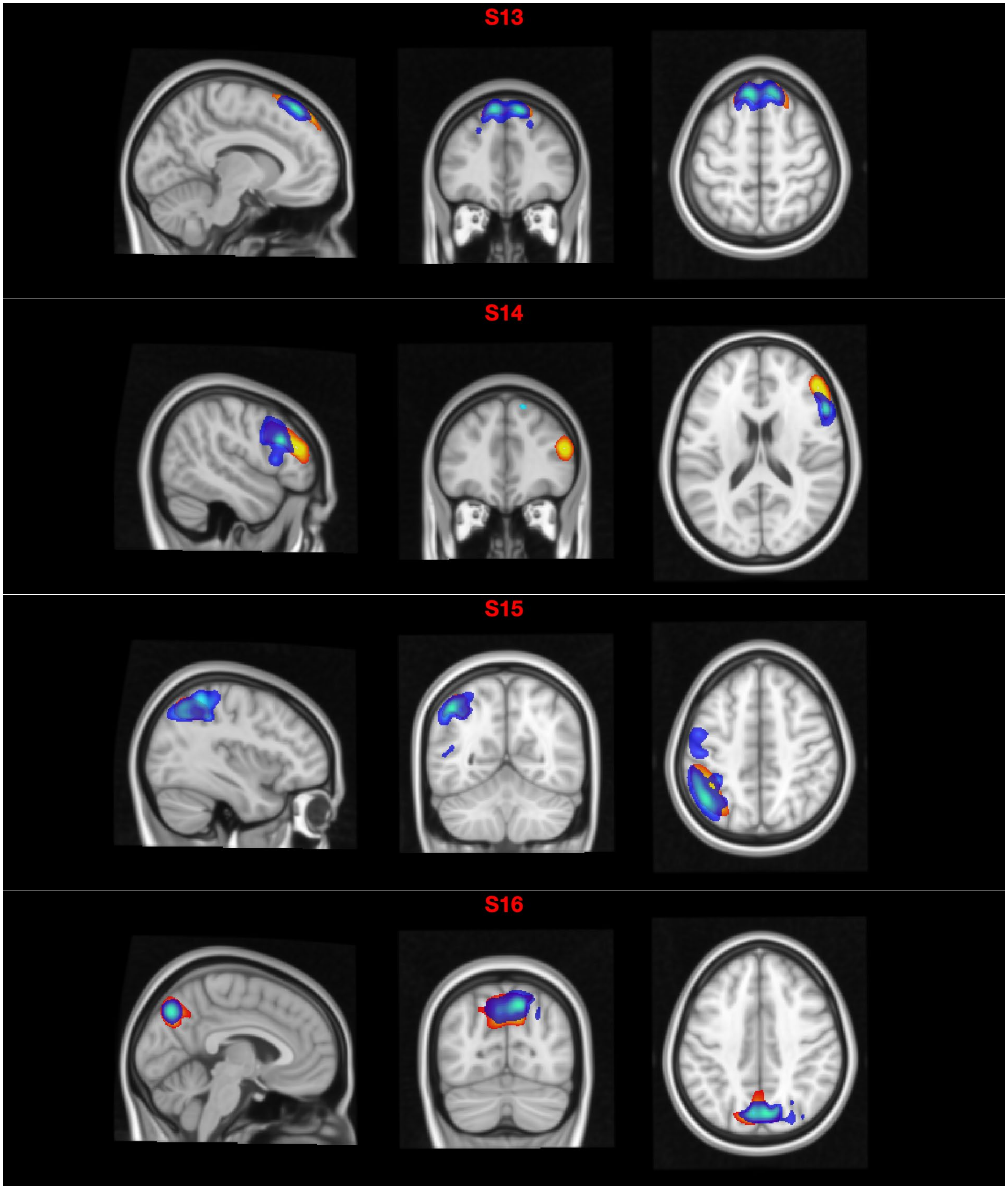
S13
S14
S15
S16

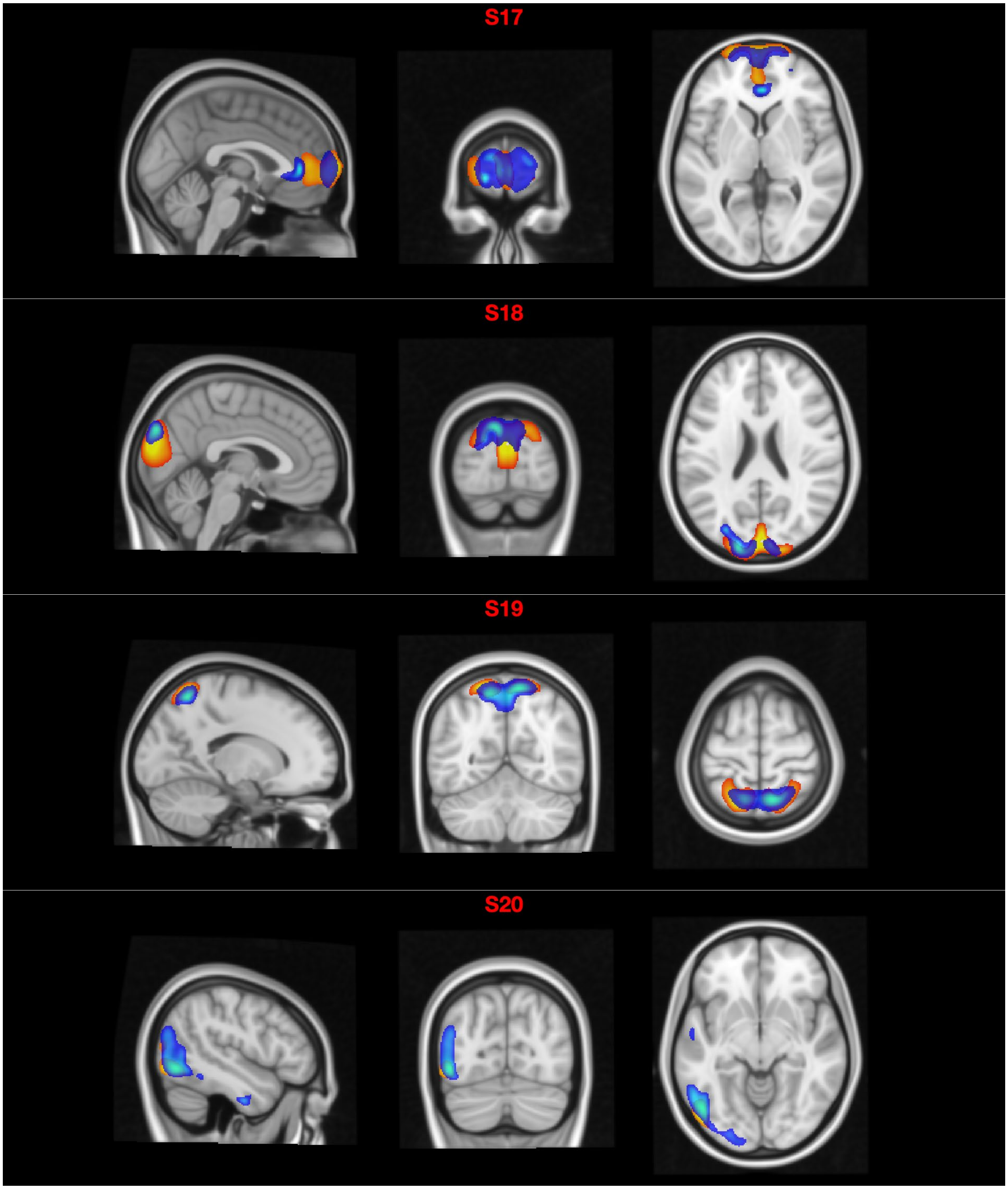
S17
S18
S19
S20

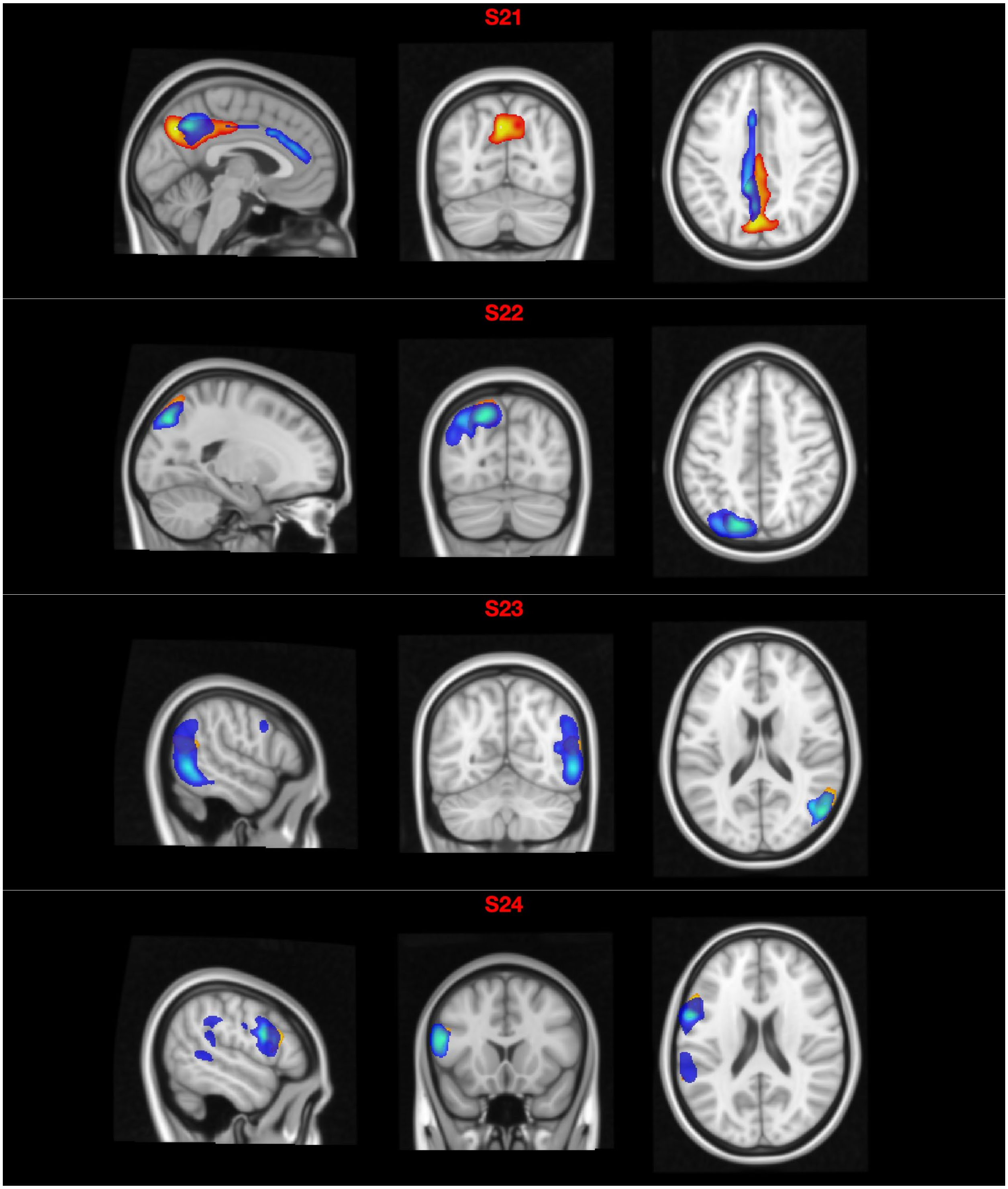
S21
S22
S23
S24

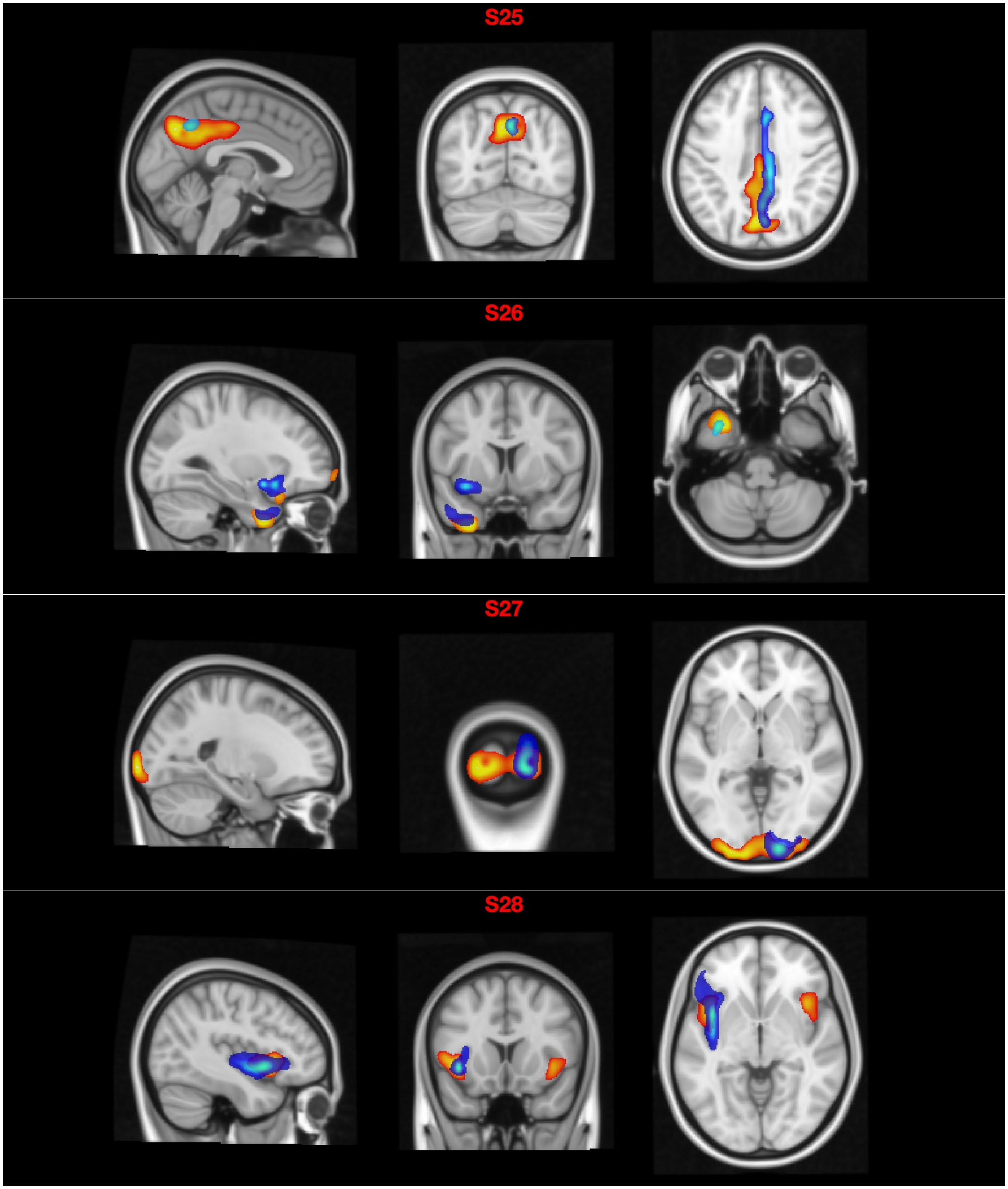
S25
S26
S27
S28

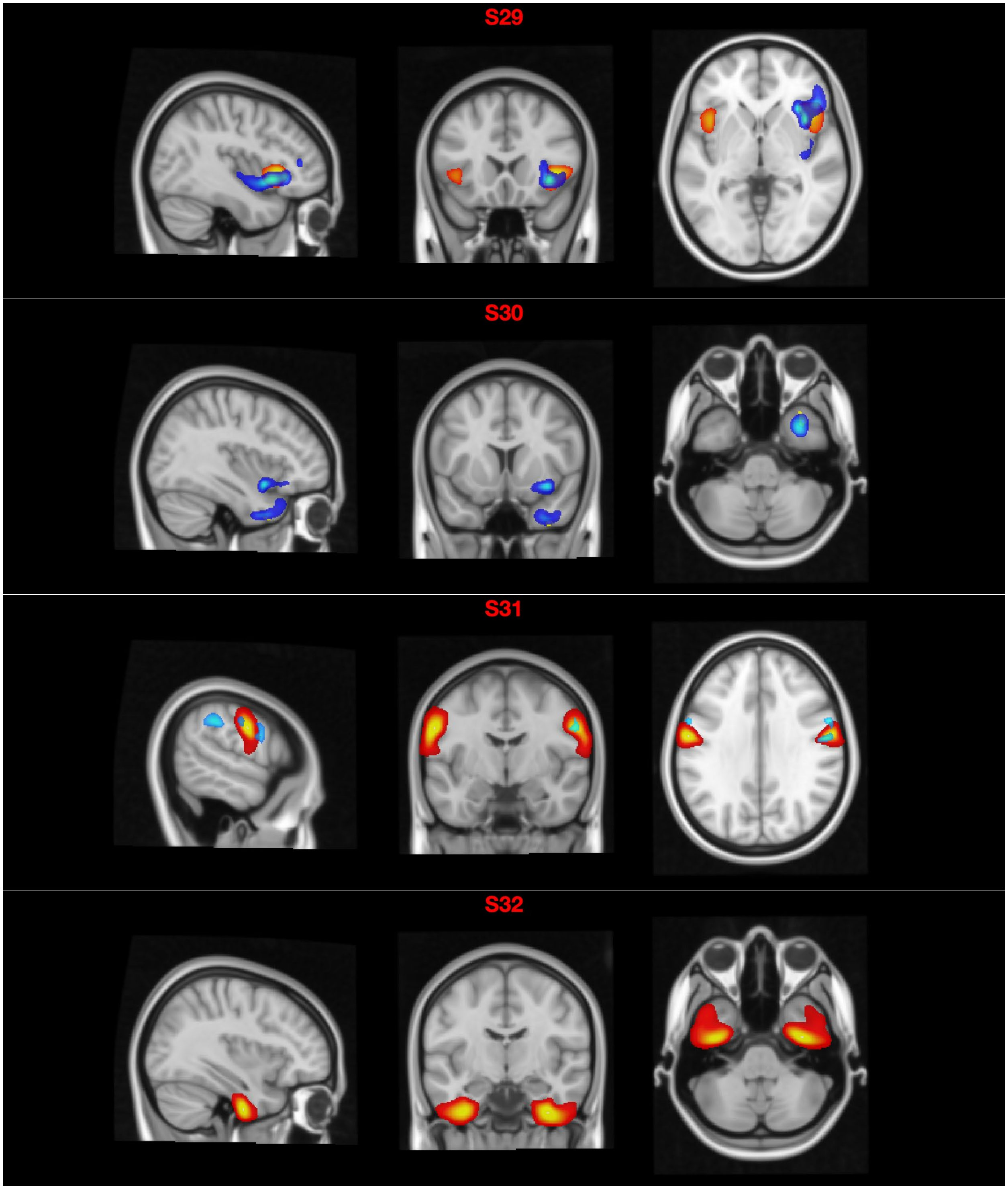
S29
S30
S31
S32

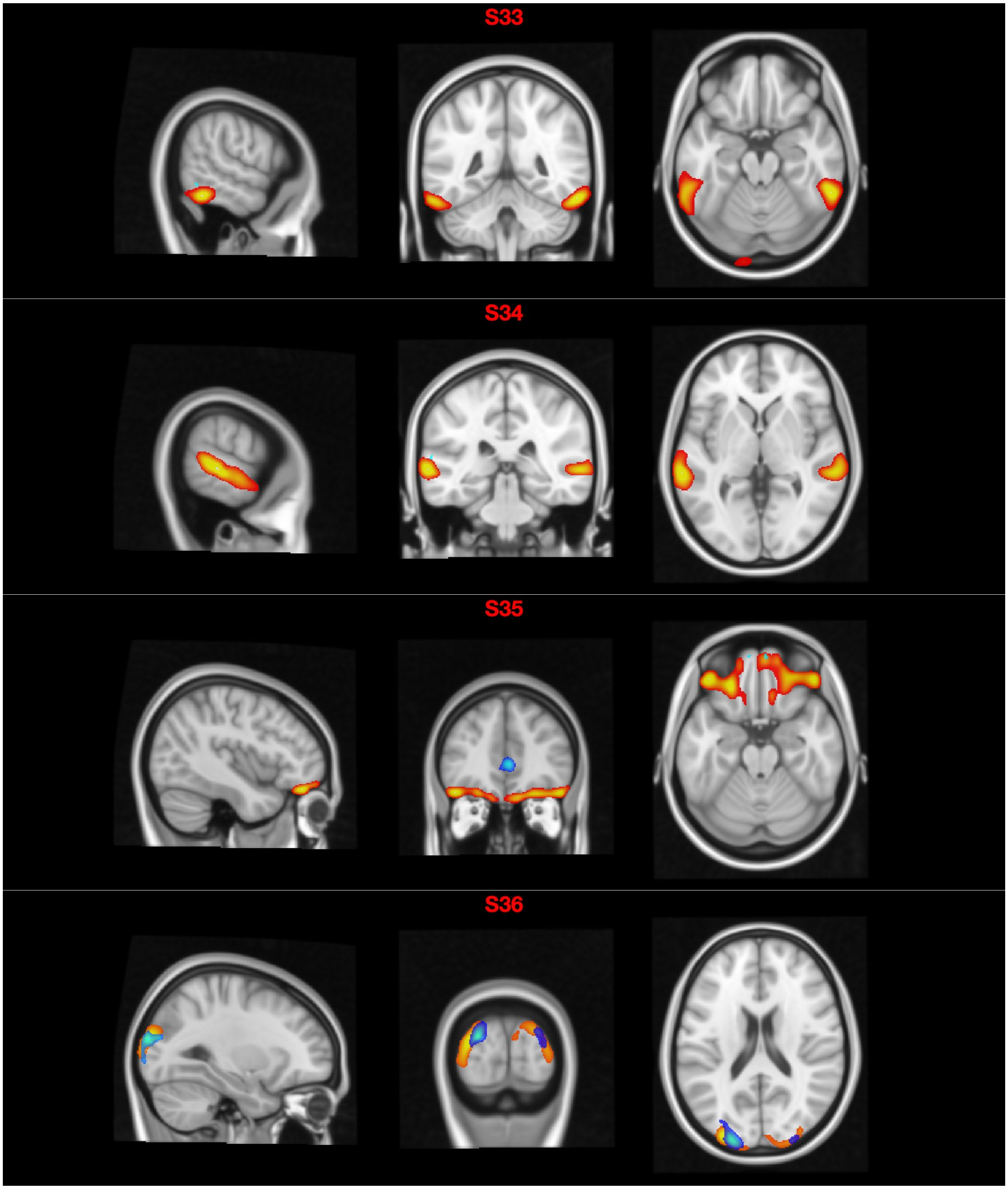
S33
S34
S35
S36

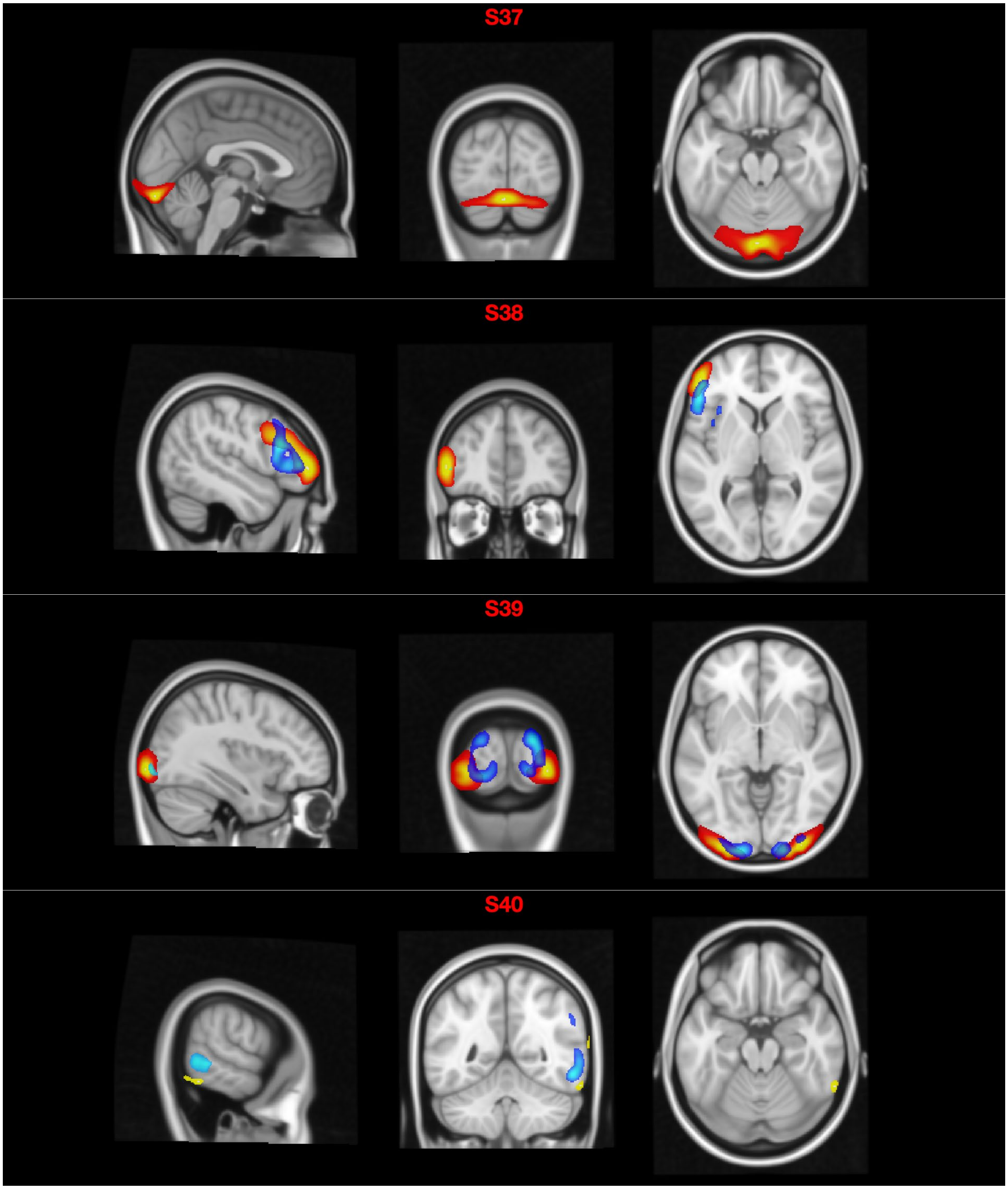
S37
S38
S39
S40

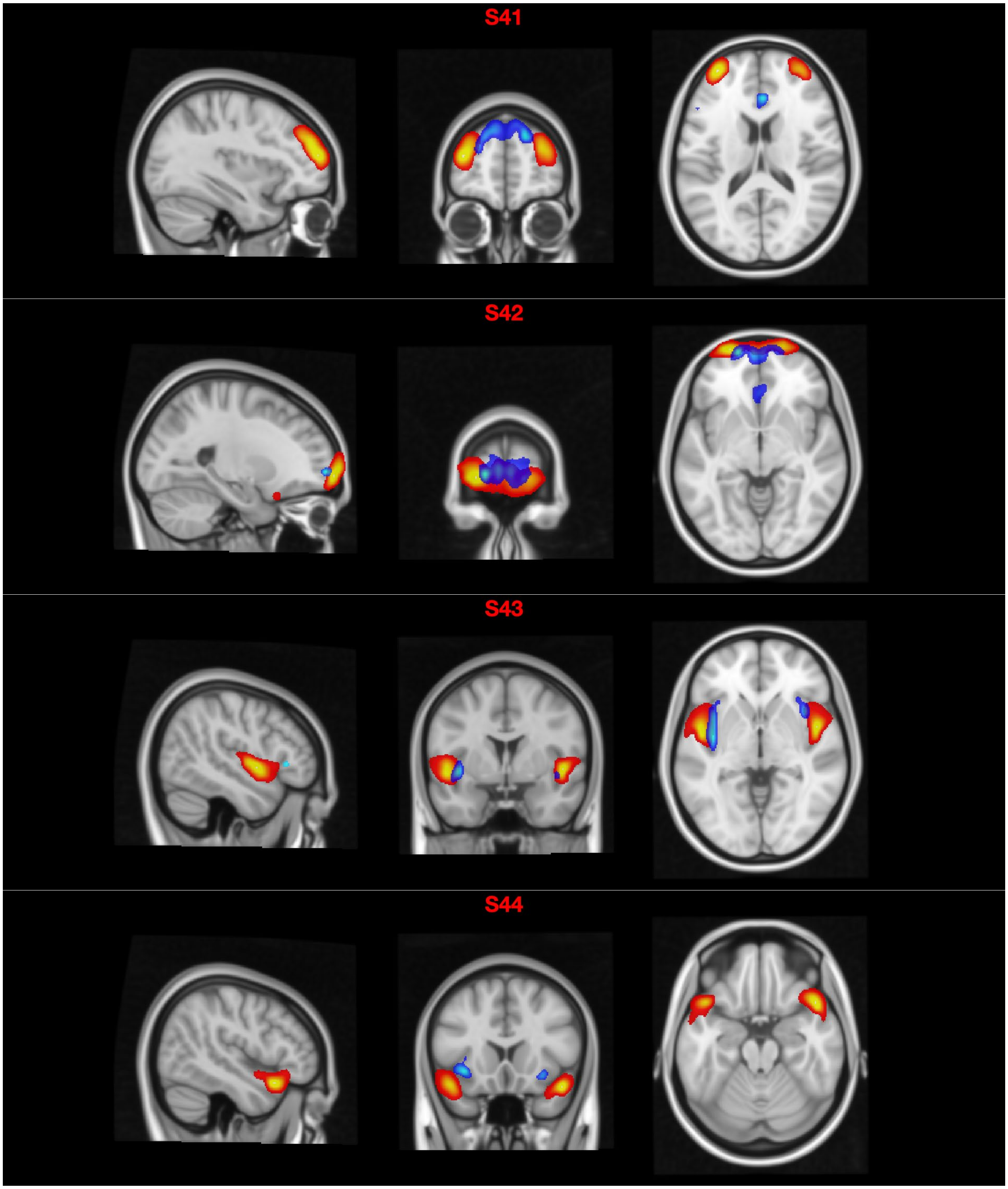
S41
S42
S43
S44

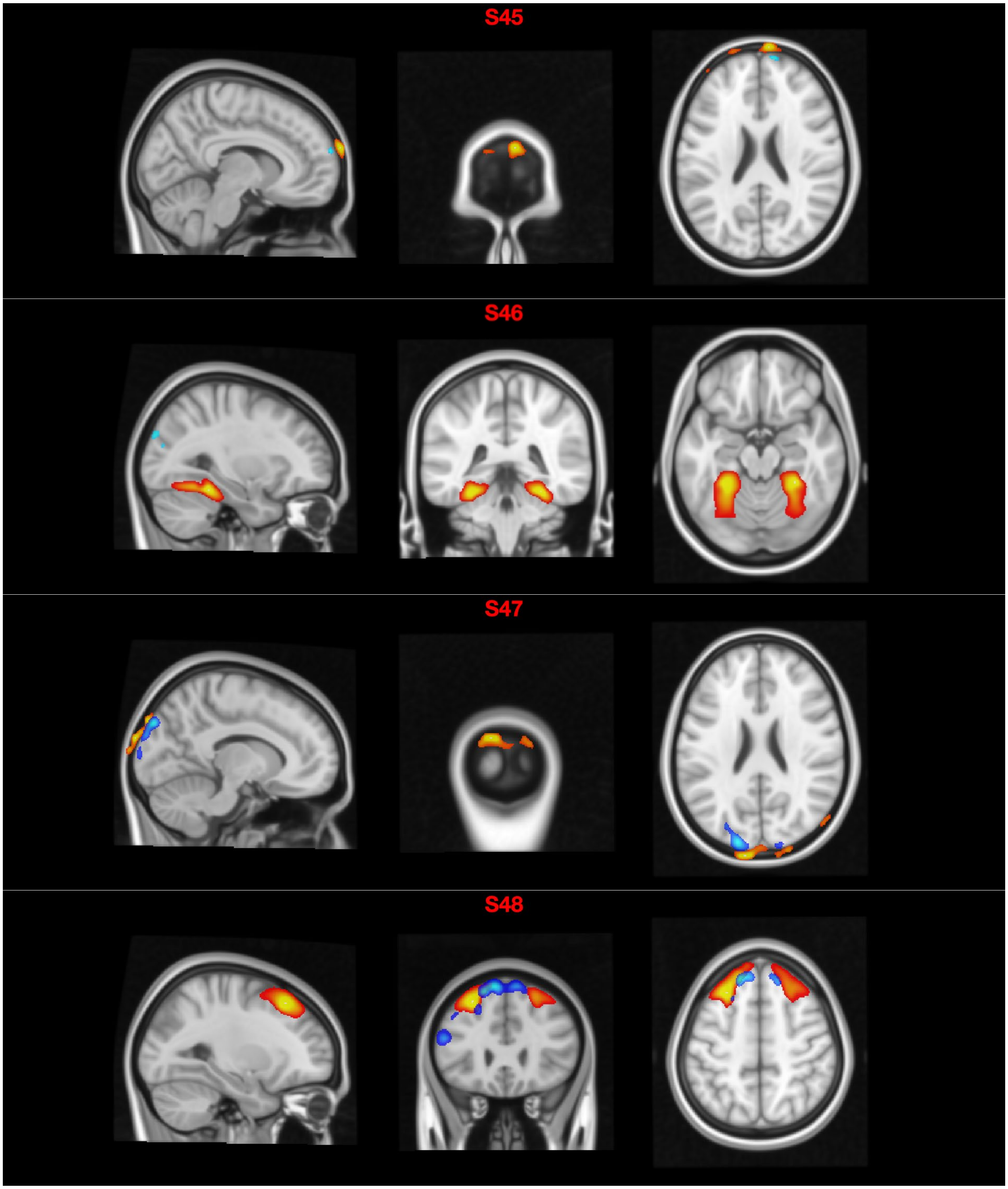
S45
S46
S47
S48

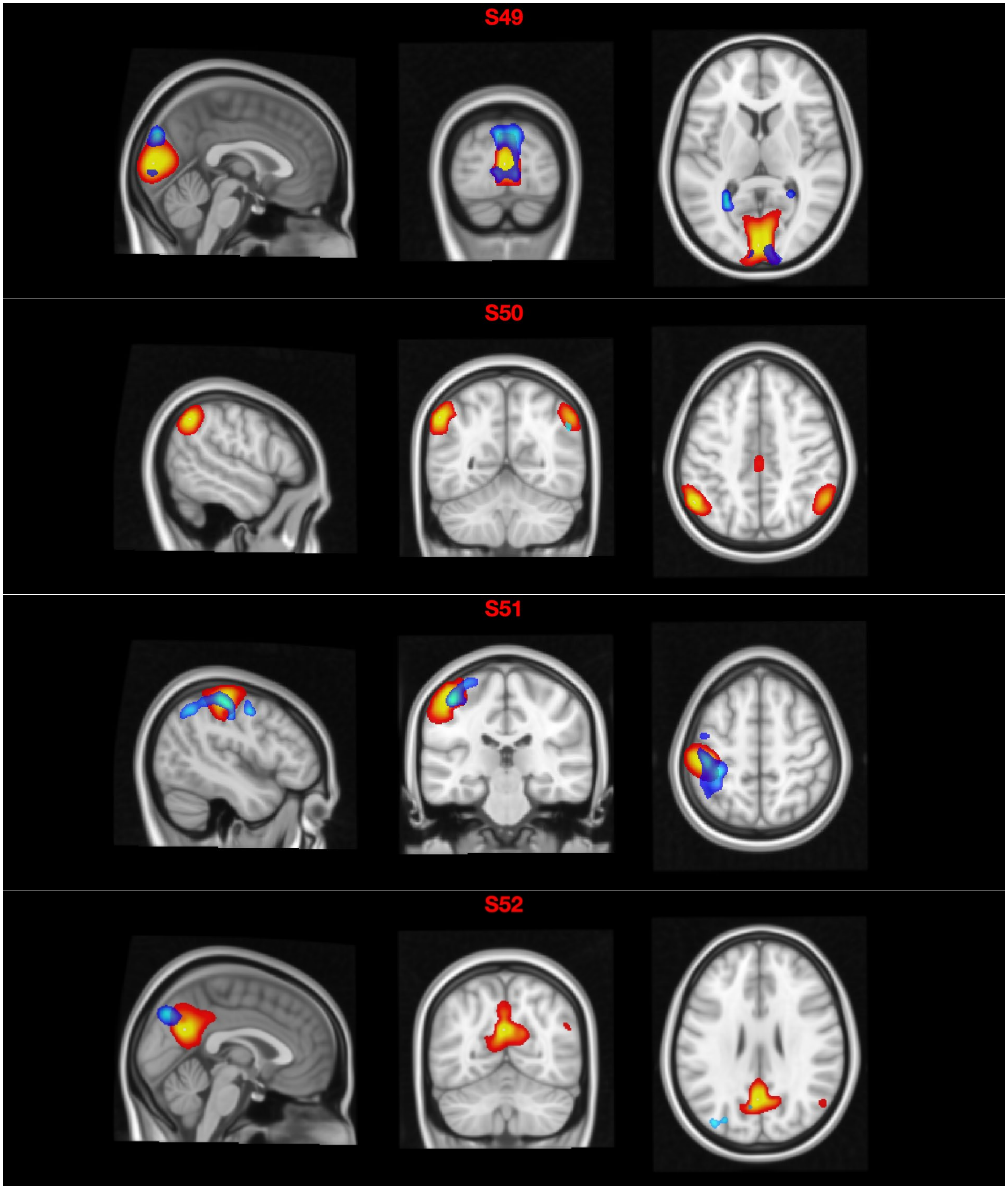
S49
S50
S51
S52

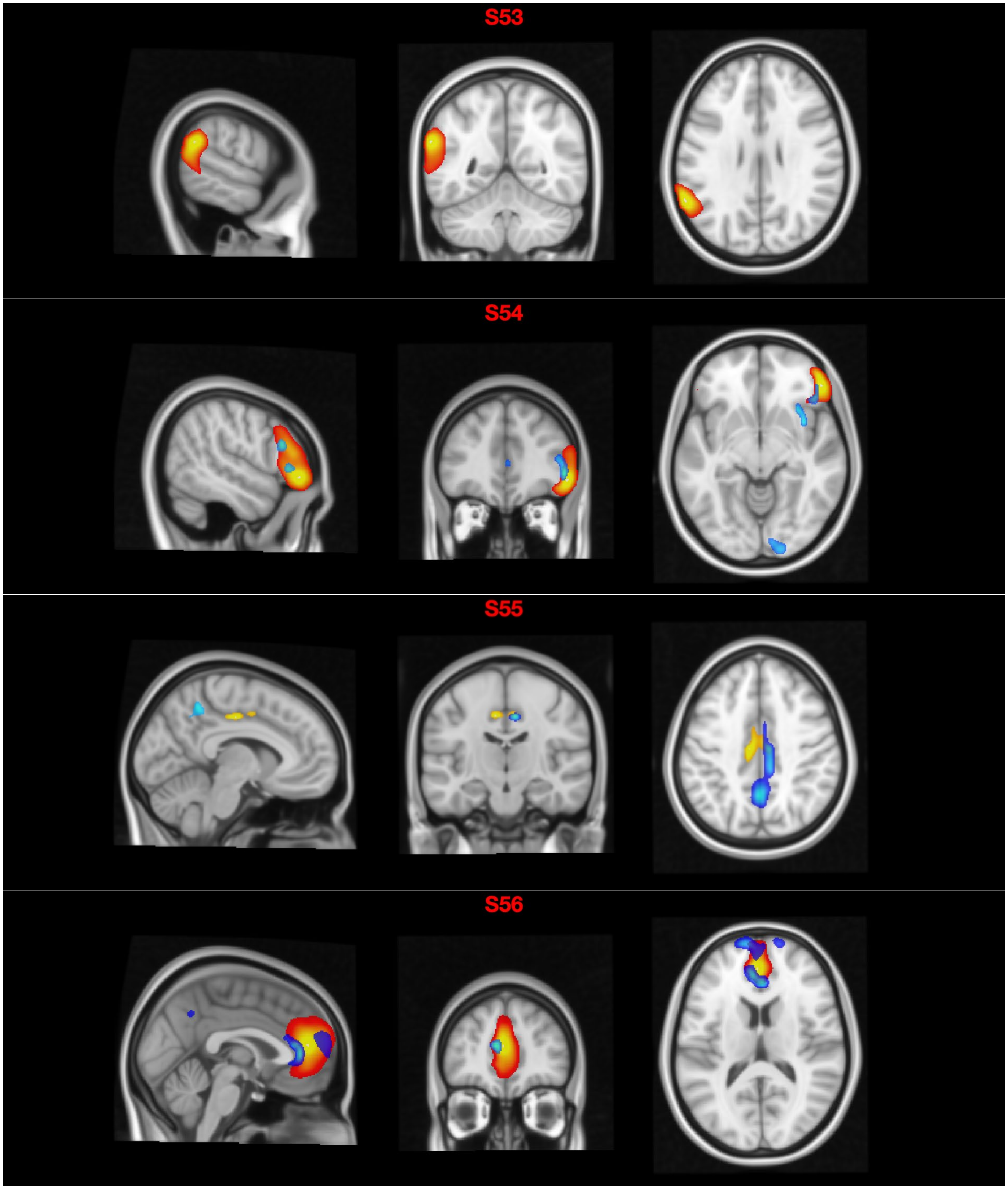
S53
S54
S55
S56

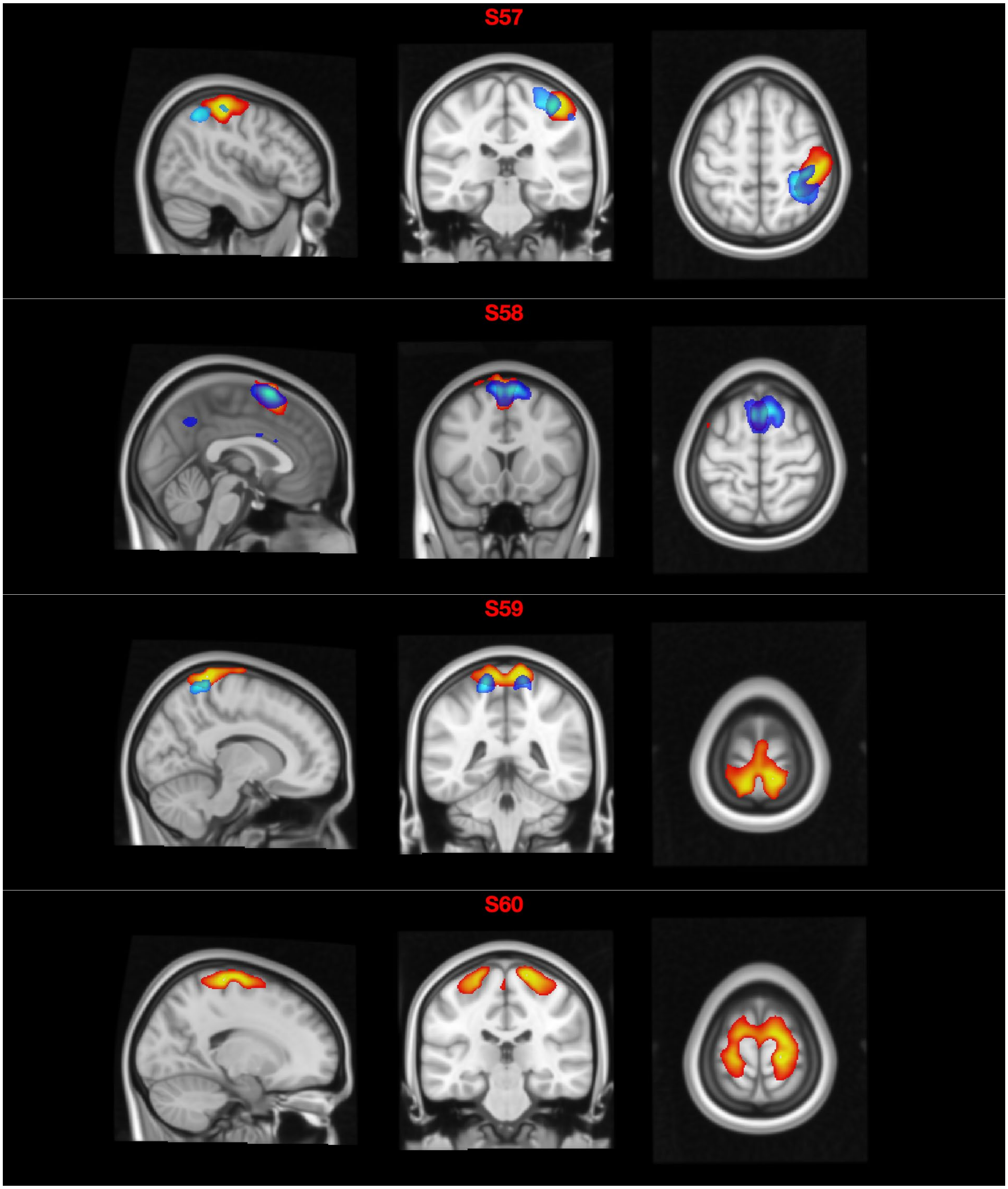
S57
S58
S59
S60

3. Joint cmICA *R* maps of FCN (red) and WMT (blue)

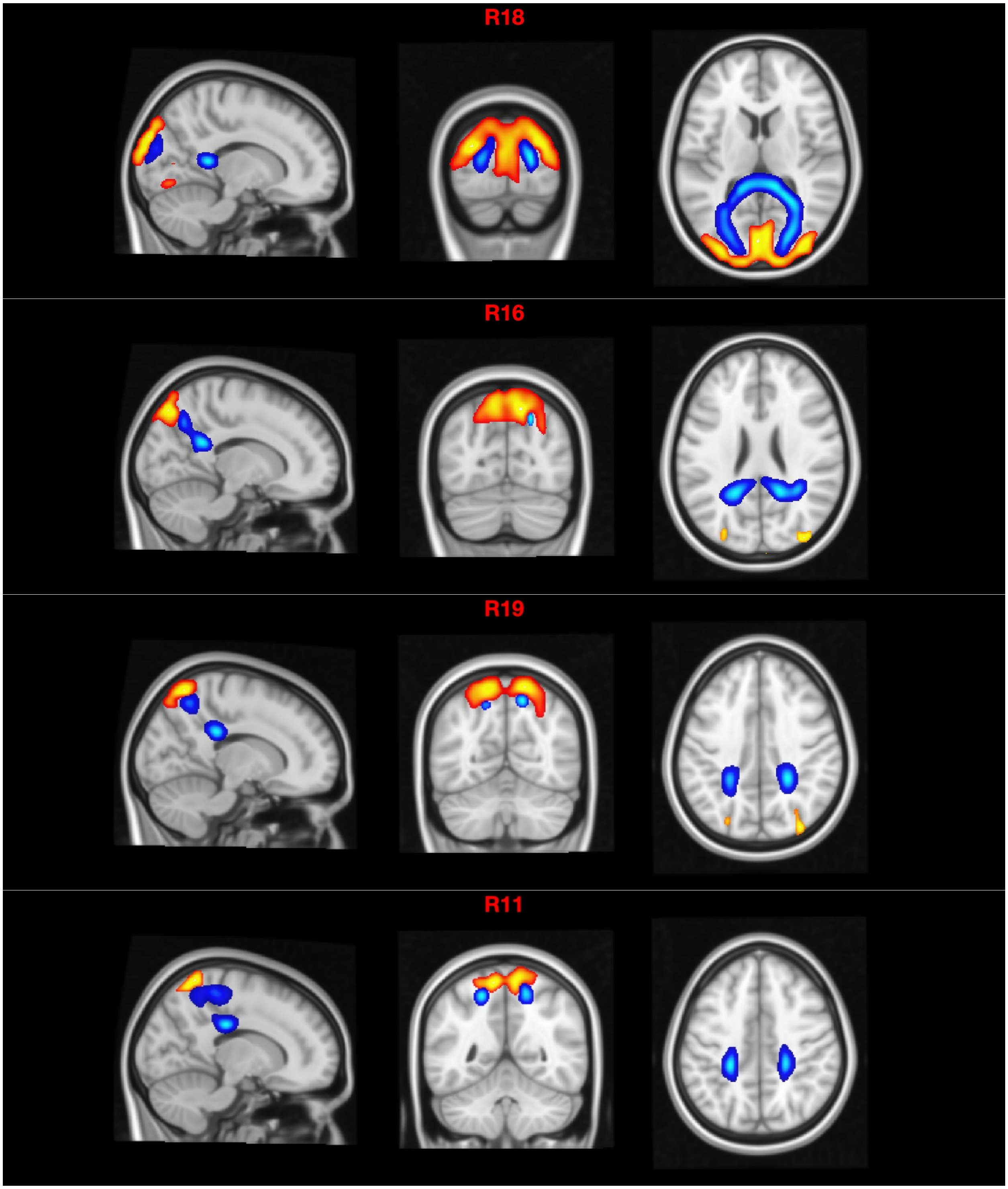

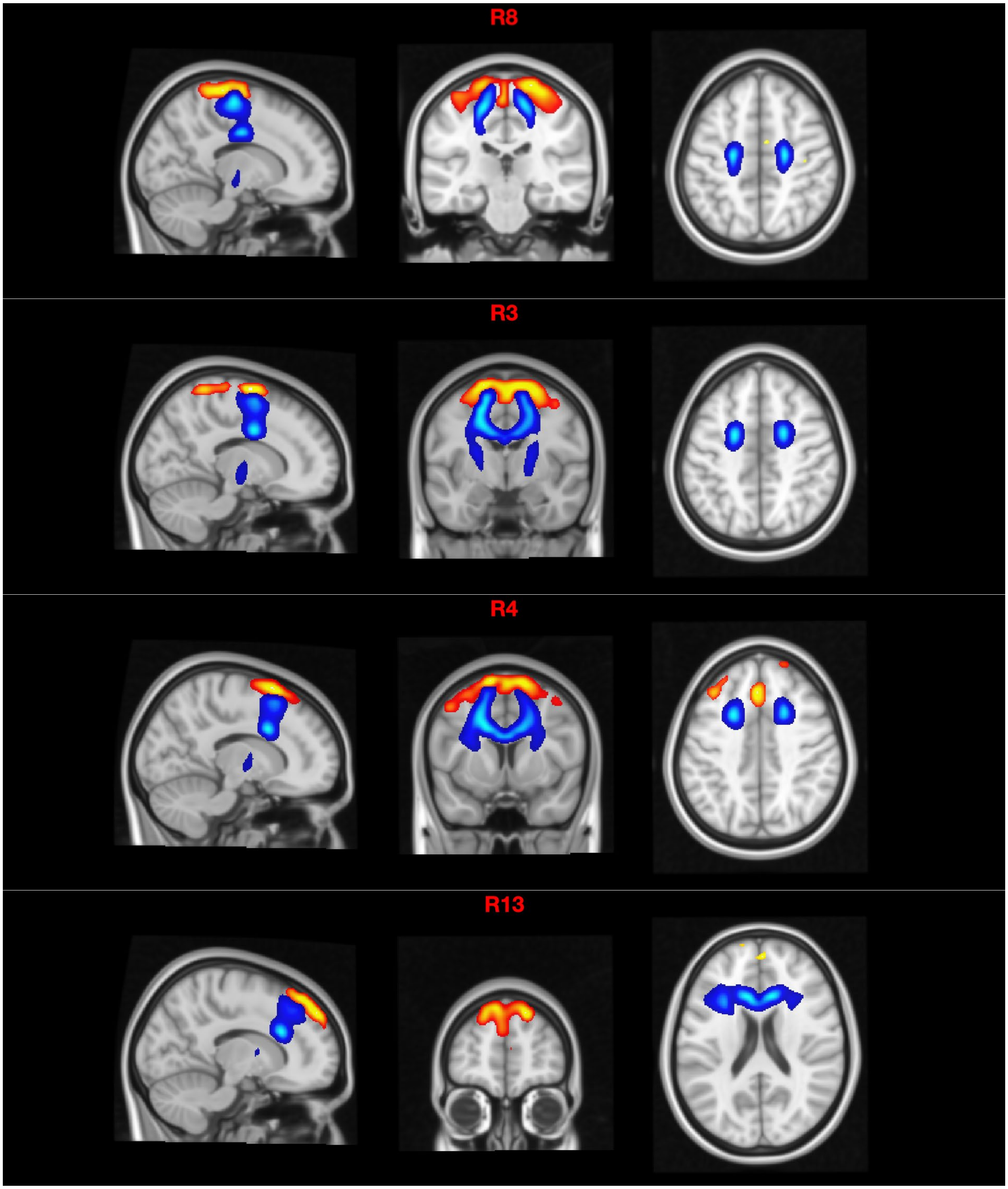
R8
R3
R4
R13

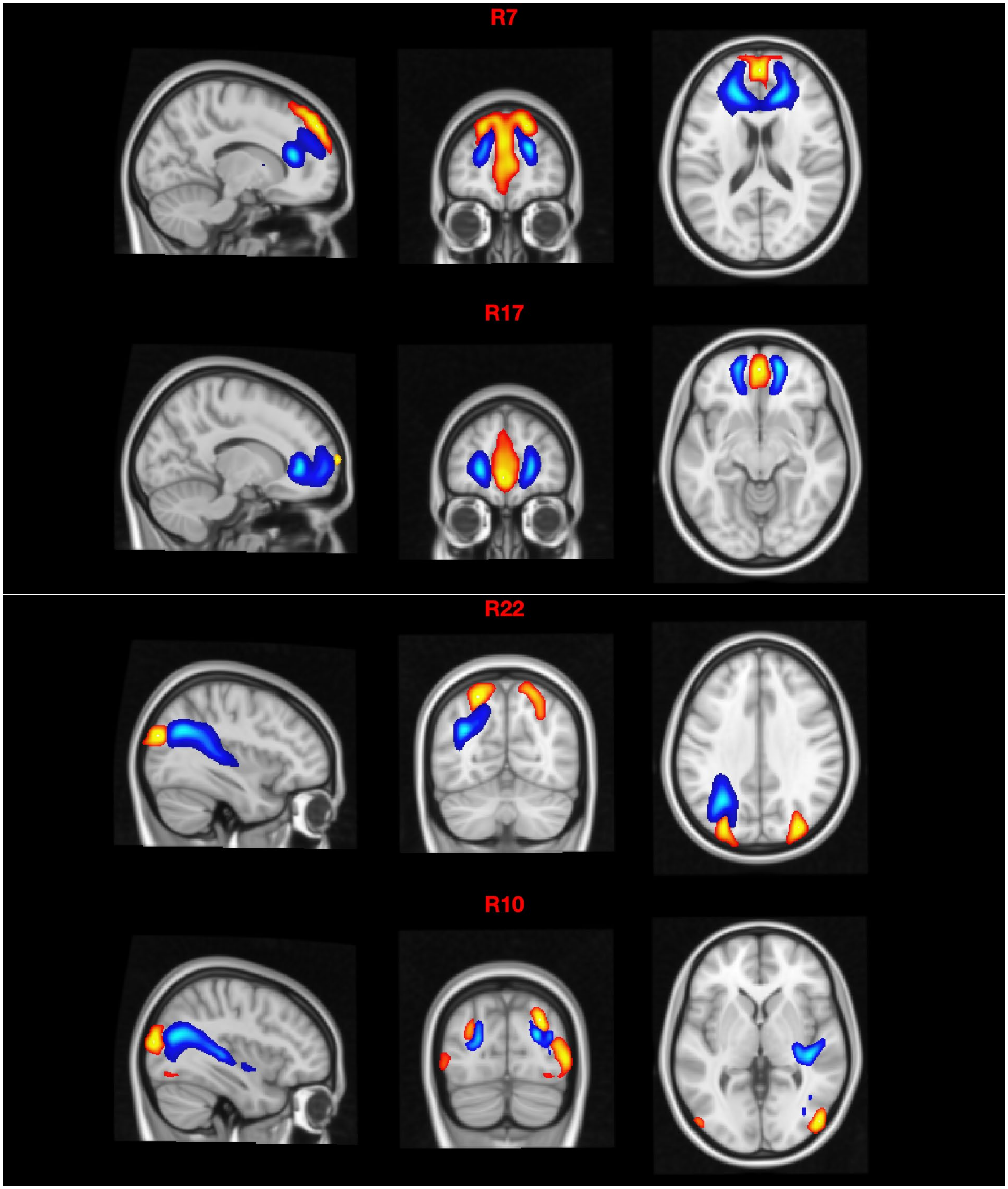
R7
R17
R22
R10

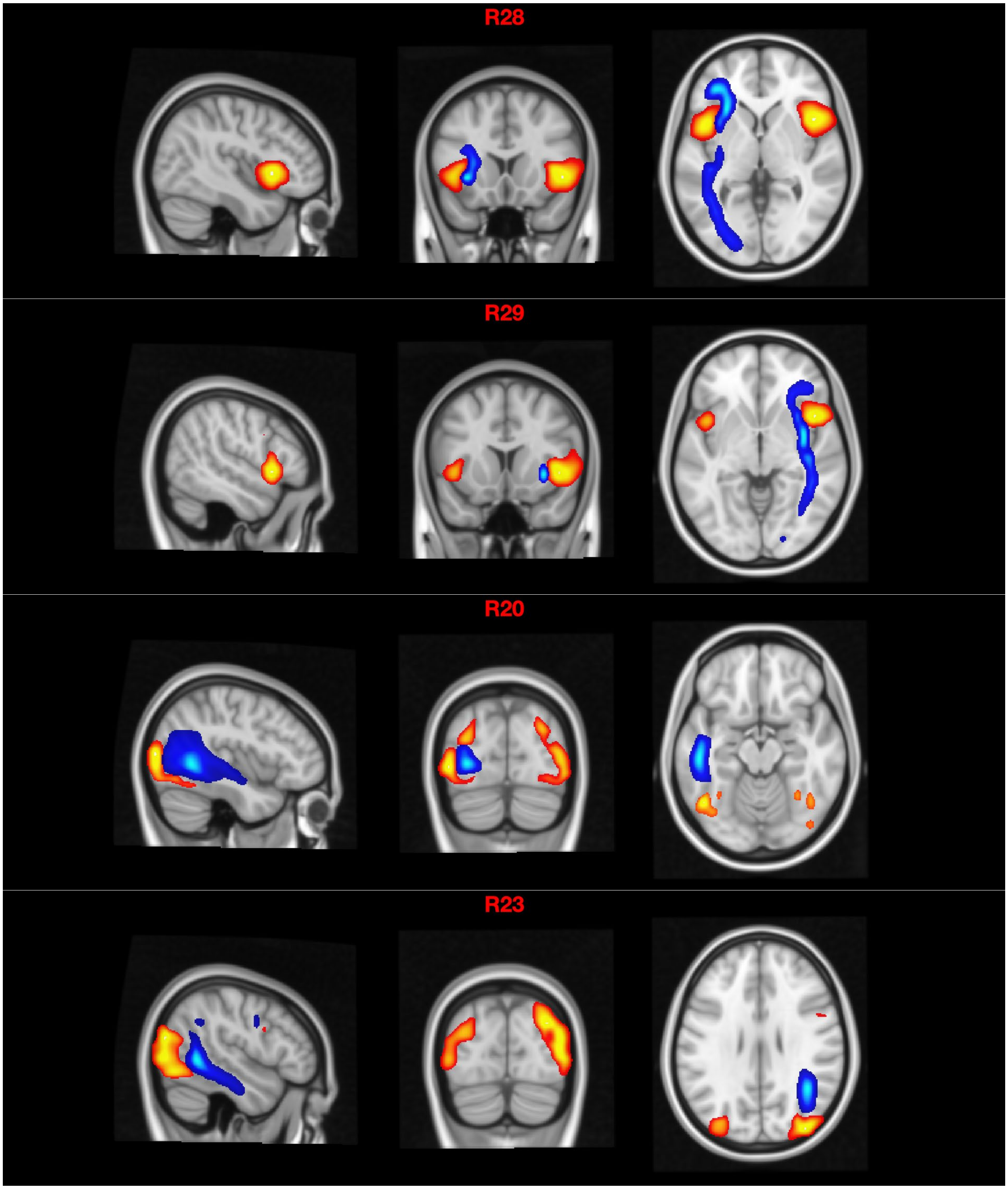
R28
R29
R20
R23

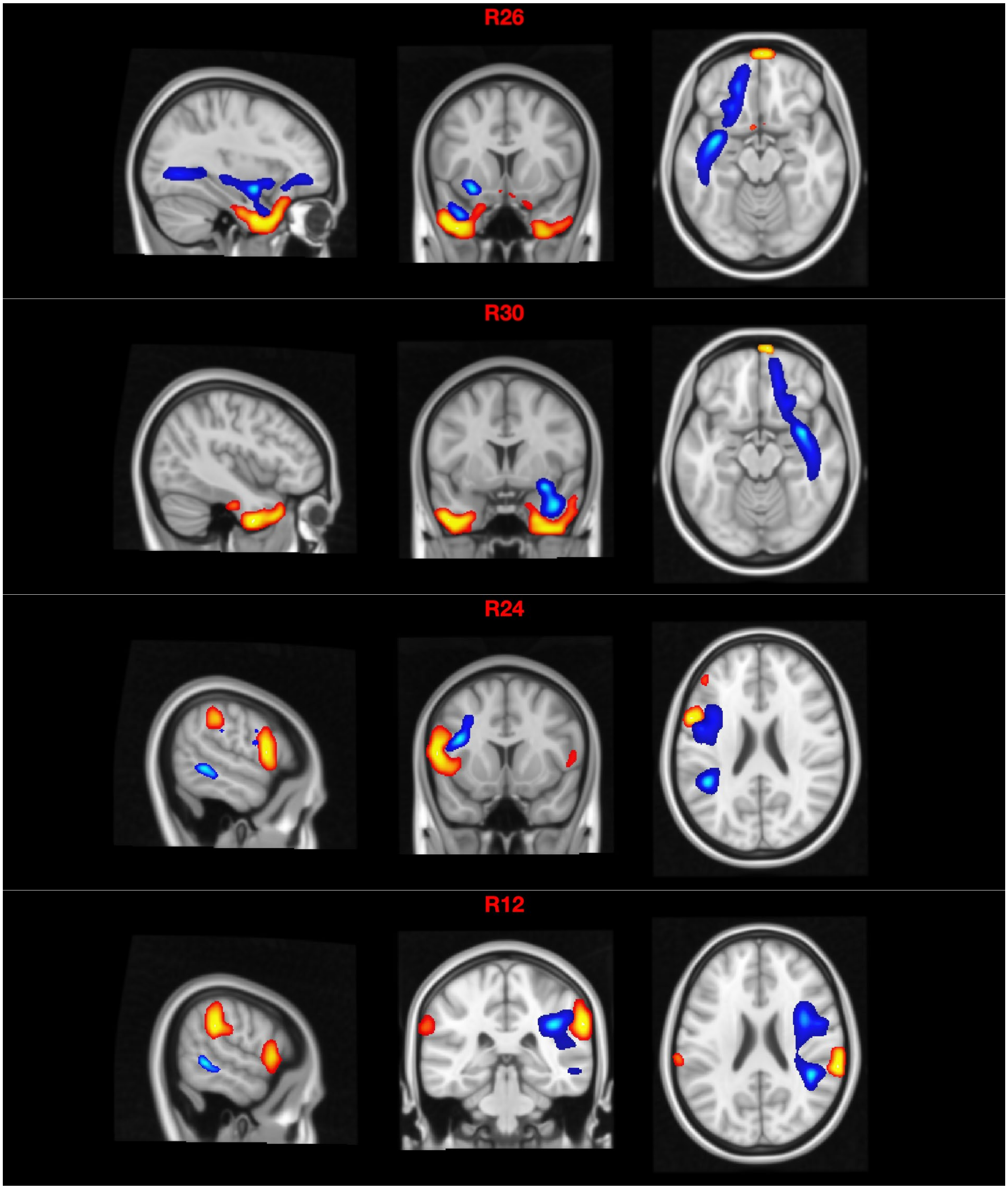
R26
R30
R24
R12

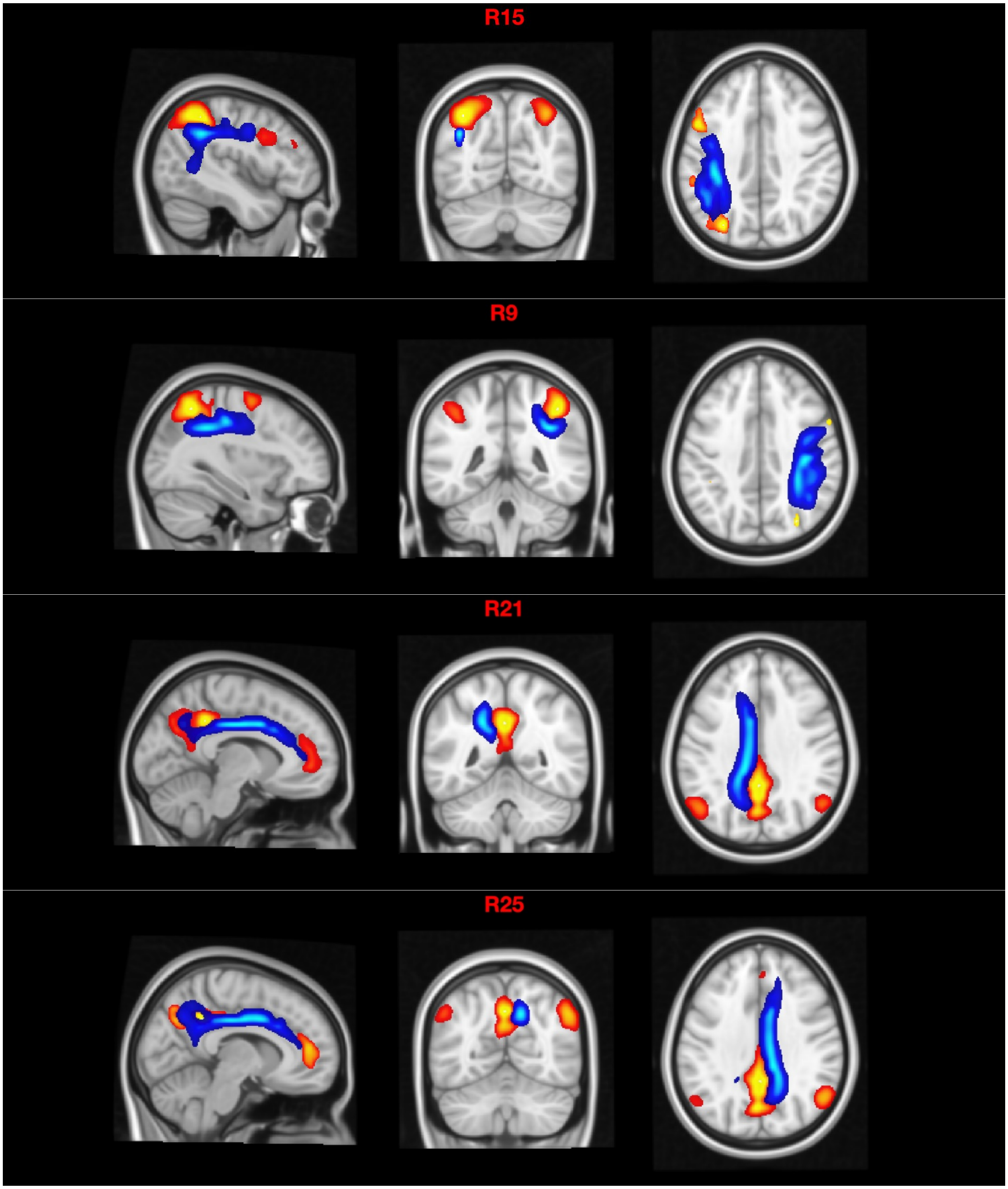
R15
R9
R21
R25

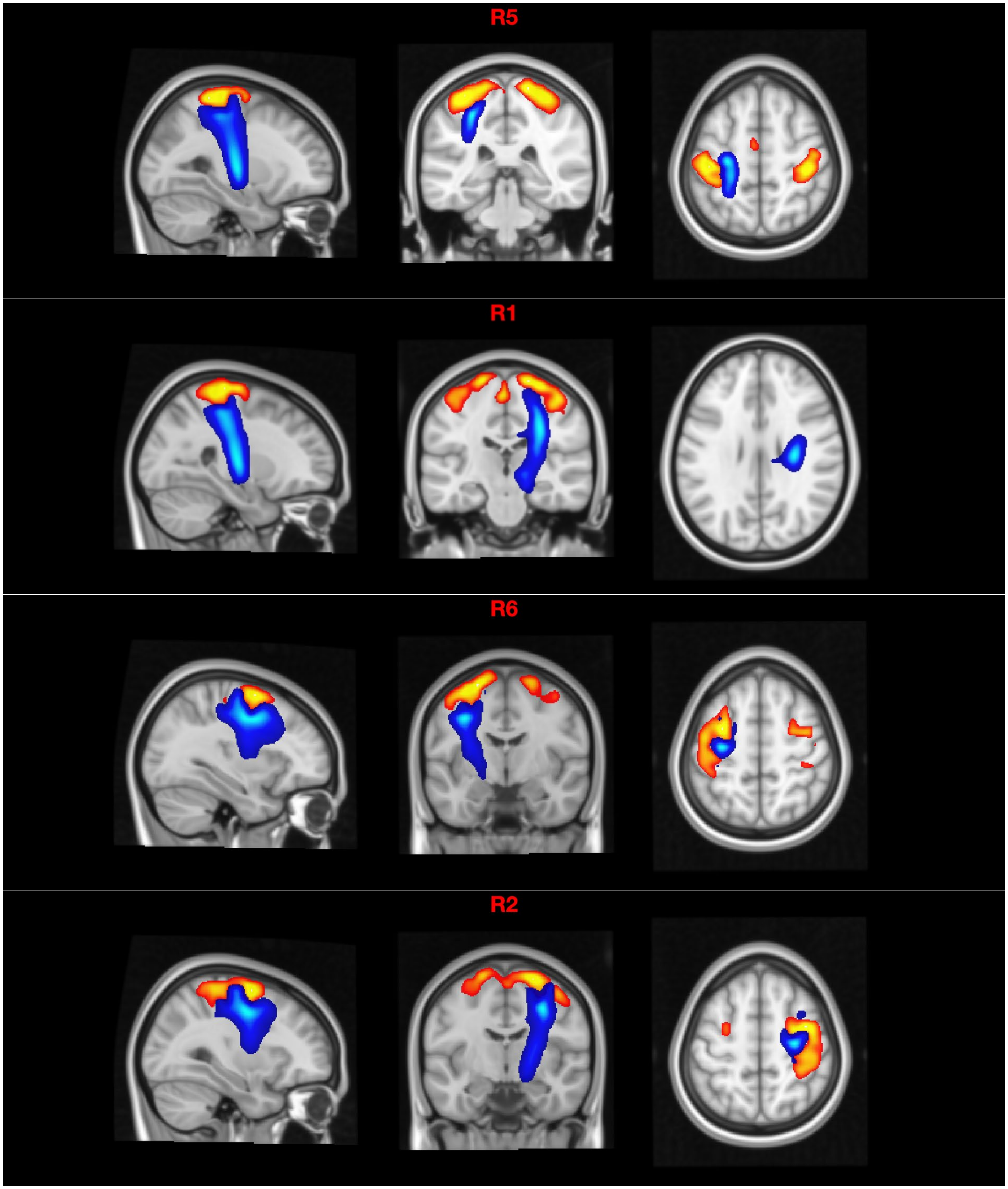
R5
R1
R6
R2

4. Source contribution and splitting

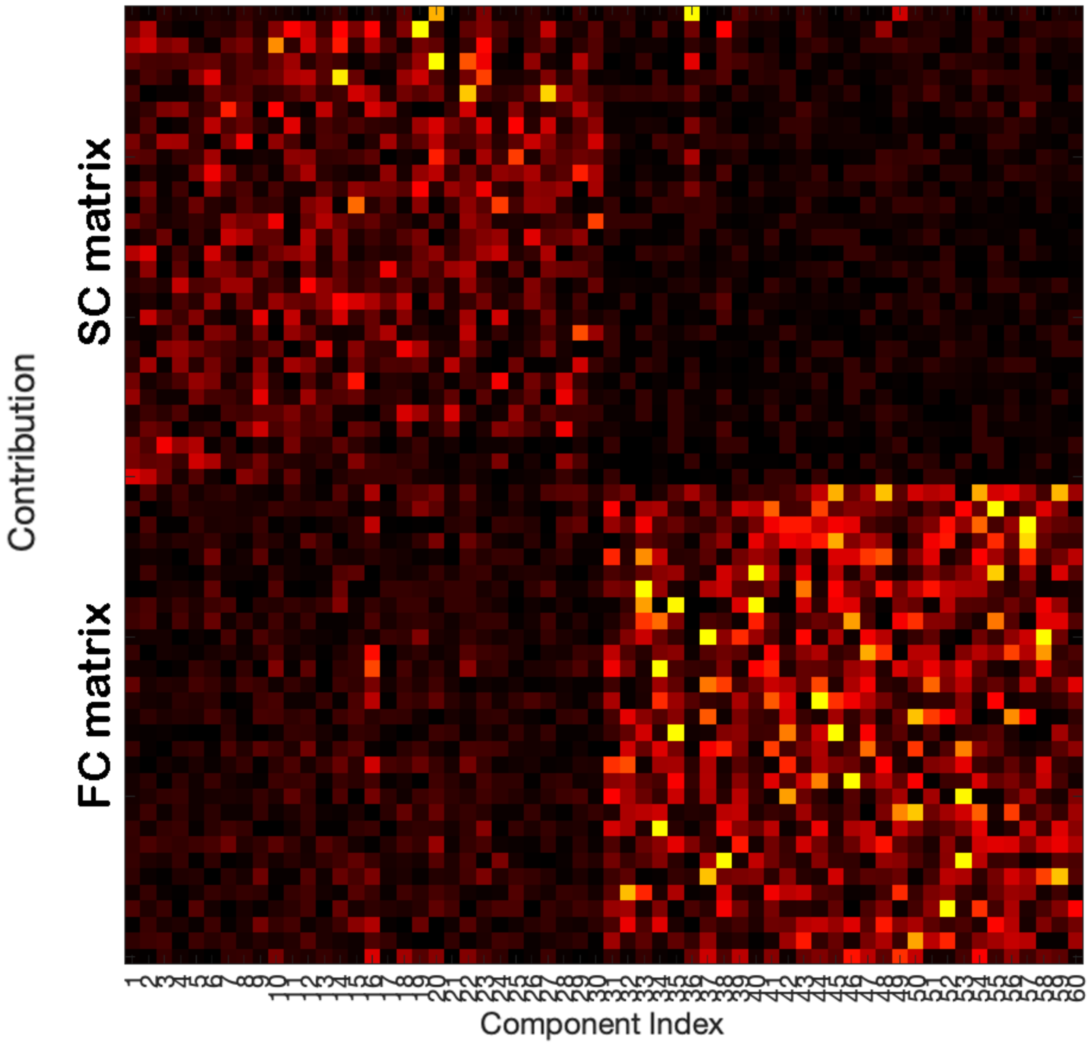
